# An *ALPK3* truncation variant causing autosomal dominant hypertrophic cardiomyopathy is partially rescued by mavacamten

**DOI:** 10.1101/2024.09.30.615779

**Authors:** Lisa Leinhos, Paul Robinson, Giulia Poloni, Sophie Broadway-Stringer, Julia Beglov, Adam B. Lokman, Gillian Douglas, Sajjad Nuthay, Oveena Fonseka, Manuel Schmid, Evie Singer, Charlotte Hooper, Kate Thomson, Richard D. Bagnall, Jodie Ingles, Christopher Semsarian, Elizabeth Ormondroyd, Christopher N. Toepfer, Benjamin Davies, Charles Redwood, Hugh Watkins, Katja Gehmlich

## Abstract

The *ALPK3* gene encodes alpha-protein kinase 3, a cardiac pseudo-kinase of unknown function. Heterozygous truncating variants (*ALPK3tv)* can cause dominant adult-onset Hypertrophic Cardiomyopathy (HCM). Here we confirm an excess of *ALPK3tv* in sarcomere-gene negative HCM patients.

Moreover, we generated a novel knock-in mouse model carrying an *ALPK3tv* (K201X). Homozygous animals displayed hypertrophy and systolic dysfunction. Heterozygous animals demonstrated no obvious baseline; however, they had an aggravated hypertrophic response upon chronic adrenergic challenge.

Isolated cardiomyocytes from heterozygous and homozygous mice showed reduced basal sarcomere length with prolonged relaxation, whilst calcium transients showed increased diastolic calcium levels and prolonged re-uptake. We also observed reduced fractional shortening. Protein kinase A-mediated phosphorylation, including that of cardiac troponin I, was significantly decreased. In agreement with the cellular HCM phenotype, reduced ratios of myosin heads in the super-relaxed state were measured. Contractile and calcium handling defects were partly corrected by treatment with mavacamten, a novel myosin inhibitor.

For the first time with a non-sarcomere HCM variant, we have demonstrated hallmark changes in cardiac contractility and calcium handling, some resembling sarcomere-positive HCM, while others are distinct and DCM-like. Mavacamten is able to partially rescue the cellular phenotype, hence could be beneficial to HCM patients with *ALPK3tv*.

## Introduction

Alpha-protein kinase 3 (ALPK3) was initially classified as an atypical protein kinase, with cardiac and skeletal muscle specific expression. The protein contains two amino-terminal immunoglobulin-like domains (Ig) and a carboxy-terminal, putative kinase domain. Based on sequence homology, it belongs to the unique family of eukaryotic alpha-protein kinases ^1^. Two studies have presented strong evidence that the protein is a pseudokinase without kinase activity ^2,3^; although another study has shown that the presence of ALPK3 regulates the phosphorylation of certain myofilament proteins; this is likely to be an indirect effect ^4^.

Early studies reported that nuclear ALPK3 plays a major role in cardiac differentiation and cardiomyocyte proliferation ^5^. More recently, the protein has been shown to localise to the M-band in mature cardiomyocytes and additionally to the nuclear envelope in differentiating induced pluripotent stem cell (iPSC) derived cardiomyocytes. In both structures, it may act as a scaffold for M-band proteins such as myomesin, and buffer force ^2^. It was also demonstrated that ALPK3 interacts with p62 within the M-band, suggesting a role in maintaining sarcomeric proteostasis ^4^.

Genetic variants in the *ALPK3* gene have been associated with cardiomyopathy ^6–10^. Recessive (homozygous) *ALPK3tv* cause an early-onset severe form of cardiomyopathy *in utero* or in early infancy, which is characterised by systolic dysfunction, dilatation and hypertrophy ^6,11,12^. Some reports also described extra-cardiac features, e.g. musculoskeletal and dysmorphic abnormalities, in the presence of recessive *ALPK3tv*. The majority of heterozygous *ALPK3tv* carriers in these families did not display an overt cardiomyopathy phenotype, even in adulthood. Nevertheless, 2 out of 10 (20%) heterozygous family members presented with adult onset hypertrophic cardiomyopathy (HCM) ^6^.

More recently, data from large cohort studies have supported the hypothesis that heterozygous *ALPK3tv* are causal of autosomal dominant HCM. Lopes *et al.* detected heterozygous *ALPK3tv* in ∼1.6% of HCM cases, and found evidence of enrichment in cases compared to the gnomAD population (odds ratio [OR] ∼16) ^9^. Evidence of enrichment of rare heterozygous *ALPK3tv* has since been described in other HCM cohorts ^8,13^. Additionally, co-segregation of a heterozygous *ALPK3tv* with autosomal dominant HCM was described in a four-generation family with seven affected members ^8^.

In this study, we undertook case control analyses to confirm the level of enrichment of rare *ALPK3tv* in our cohort of HCM patients. To examine the functional consequences, we generated a novel knock-in mouse model carrying a heterozygous *ALPK3tv* (K201X) found in an adult onset autosomal dominant HCM patient. While the heterozygous mice showed no overt phenotype *in vivo*, pathological dysfunction that shared features seen with sarcomere-related HCM was evident in isolated cardiomyocytes. We also demonstrated that mavacamten, a drug licenced to treat patients with obstructive HCM, can revert some aspects of the cellular dysfunction observed in cardiomyocytes of our mouse model.

## Material and Methods

### Ethical statement: patient data

All research involving humans meets the ethical standards of the 1964 Declaration of Helsinki and its amendments. Patients provided informed consent for genomic sequencing, and analysis of demographic, clinical, and family history data through the NIHR Bioresource Rare Disease HCM project (BRRD, Research Ethics Committee reference 13/EE/0325) or earlier studies of the genetic basis of HCM.

### Patient case-control analysis

Rare variant burden analyses were performed as described previously ^14^, using rare variant data available from 230 unrelated, sarcomere variant negative, HCM cases and 6,219 unrelated individuals, recruited to other rare disease projects within the BRRD. Well-established sarcomere-related HCM genes (*MYBPC3, MYH7, TNNI3, TNNT2, MYL2, MYL3, ACTC1, TPM1*), genes for common differential diagnoses (*PRKAG2, GLA, FHL1*), and other more rarely associated, but validated, HCM genes (*CSRP3, PLN*) were included in the genetic testing.

The proportion of rare (total minor allelic frequency <1x10^-4^ in gnomADvs2.1.1) missense, truncating (frameshift, nonsense, splice donor/acceptors), non-truncating (missense, in-frame insertions and deletions) and synonymous *ALPK3* variants was compared between HCM cases and controls. Fisher’s exact test (FET) and odds ratios (OR) with 95% confidence intervals (CI) were calculated.

Variants were annotated according to the MANE transcripts (NM_929778.5; ENST0000025888.6). Variants were reviewed and classified according to clinical variant interpretation guidelines ^15^.

### Ethical Statement: animal procedures

All animal procedures meet the ethical standards of the 1964 Declaration of Helsinki and its amendments. Experimental animal studies were performed in accordance with the UK Home Office guidelines and approved by institutional ethical review board (Project Licence PPL PDCE16CB0).

### Generation of mouse model

A novel knock-in (KI) mouse model carrying one of the *ALPK3tv* found in a patient with autosomal dominant HCM (*ALPK3* p. K201X), was generated by the Genome Engineering Core Facility at the Wellcome Centre for Human Genetics.

To incorporate the desired c.601A>T variant in a mouse model (and two further silent changes, Fig. S1A), a CRISPR/Cas9 nuclease was designed against murine *Alpk3* exon 5 (ENSMUSE00000506542) the CRISPOR design tool (crispor.telfor.net) along with a silent mutation to remove the PAM sequence adjacent to the target sequence. For easy detection of the recombinant allele, an additional silent mutation was introduced, resolving in a unique HindIII restriction site. One target site was identified close to the Lysine residue according to genome-wide specificity (lack of many significant off-target sites). A sgRNA designed against the selected guide sequence (5’- GCATGGAAAAGAGGCTTCAG-3’) was purchased from Synthego. A founder mouse was generated by microinjecting the 139 nt ssODN and the guide-RNA for the designed CRISPR/Cas9 nuclease into fertilized C57BL/6J zygotes.

Animals were backcrossed onto C57BL/6JOlaHsd (Envigo RMS (UK) Ltd) for at least 6 generations until congenic before phenotyping. Consistent genetic background of mice was confirmed using MiniMUGA genotyping array analysis via Transnetyx® service ^16^.

Genotyping was performed using the REDExtract-N-Amp™ Tissue PCR Kit (Sigma) and primer pair Alpk3-F1 5’- AGA AGA TGC TGC CAT CTA CCA A-3’ and Alpk3-R1 5’- TGA CCT CGC AGA TGT ATG TCA G-3’. The 470 bp amplicon was digested with *HindIII* restriction enzyme (NEB) and analysed by agarose electrophoresis (1.5 % agarose in Tris-Borate-EDTA buffer, Fisher). The wildtype allele was not digested (470 bp fragment), while the *Alpk3* K201X allele was digested into two fragments of 183 bp and 287 bp.

### Animal husbandry

Animals were maintained in standardised pathogen-free ventilated cages with food and water supply *ad libitum* and a 13 h light / 11 h dark circadian rhythm (150–200 lux cool white LED light, measured at the cage floor), with the only reported positives on health screening over the entire time course of these studies being for*. Chilomastix Sp* and *Entamoeba muris*. Where possible, animals were housed in social groups of mixed genotypes, except for animals under adrenergic challenge studies, which were single-housed with additional enrichment for refinement reasons.

All studies were carried out using adult, male mice littermates, while females were used for breeding. Mice were housed in mixed genotype cages. Treatment groups were assigned at random and all data acquisition and analysis was performed blinded. Animals were culled by cervical dislocation followed by confirmation of death via cessation of circulation.

### Echocardiography

Ultrasound *in vivo* phenotyping of the heart was performed on male mice under general anaesthesia using the Vevo 3100 micro-ultrasound system (FUJIFILM VisualSonics). Anaesthesia was induced by an induction chamber filled with 4% isoflurane in 2 L/min 100% oxygen and later maintained with 1.5% - 2% isoflurane (Zoetis) in 1 L/min 100% oxygen delivered through a fitted nose-cone. The animal’s body temperature was maintained at 37 °C using a heated platform during the procedure and vital functions were monitored via an embedded electrode system. Cardiac structure and function were measured at comparable physiologically relevant heart rates (where possible between 460 to 480 bpm) in 30 to max. 60 minutes, according to the approved PPL guidelines. Parameters were analysed and calculated using the Vevo LAB software (version 5.6.1).

### Adrenergic challenge via osmotic minipumps

To induce hypertrophic pathways and to explore consequences on cardiac performance, chronic adrenergic challenge with isoprenaline/phenylephrine (Iso/PE) delivered by osmotic minipumps was performed. Male animals were assigned to treatment groups at random and underwent a pre-implant baseline echocardiography.

Baseline echocardiography data were also used as additional values for the for the 3-month-old WT and Hom *Alpk3* K201X comparison (n = 23 WT, n = 8 Hom, Fig. 1, Table S3) and for the 3-month-old WT and Het *Alpk3* K201X comparison (n = 23 WT, n = 21 Het, n = 18 Hom, Fig. S2, Table S4).

**Figure 1.**
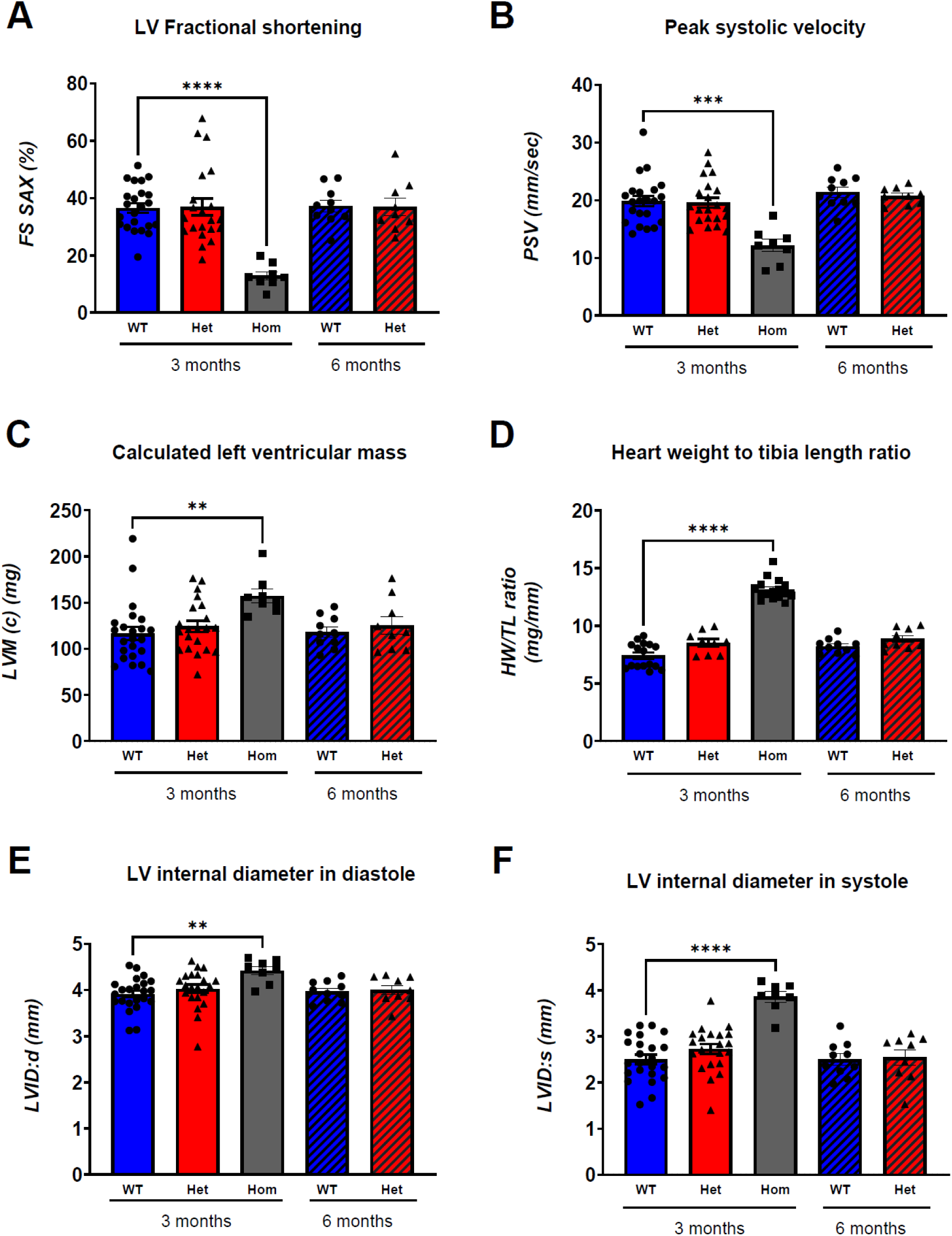
Phenotype of the novel *Alpk3 K201X* mouse model. Cardiac structure and function were assessed via echocardiography in wildtype (WT), heterozygous (Het) and homozygous (Hom) *Alpk3* K201X animals at 3 months. WT and Het animals were also assessed at 6 months. Fractional shortening (FS, **A**), peak systolic velocity (PSV, **B**), calculated left ventricular mass (LVM, **C**), heart weight (HW) to tibia length (TL) ratio (**D**), and left ventricular internal diameter in diastole (LVID:d, **E**) and systole (LVID:s, **F**) are shown. Values are presented as mean ± SEM. Kruskal-Wallis test indicates significant changes in Hom mice; ** p < 0.01, **** p < 0.0001 versus WT 3 months. To compare WT and Het mice at 6 months, unpaired Student’s t-test was performed and no significant changes were observed. For a full set of echocardiographic parameters and n numbers per parameter see Table S3.

All animals received an injection of buprenorphine (0.05 mg/kg body weight of 0.3 mg/mLVetergesic® Ceva) subcutaneously for pain prevention before the surgical procedure. Osmotic minipumps (ALZET 1002) for drug administration were implanted subcutaneously in a sterile midscapular incision procedure under general anaesthesia (constant supply of 2% isoflurane in 1 L/min 100% oxygen, as described above).

Treatment groups received isoprenaline and phenylephrine (Iso/PE; in hydrochloride forms, Sigma Aldrich), which are α- and β-adrenergic agonists, at a concentration of 15 mg kg^−1^ body weight each in 0.9% NaCl (Vetivex ® 1, Dechra) at a constant flow rate of 0.25 µl/hr per day for 14 days in total. Control groups received 0.9% NaCl as a placebo via osmotic minipumps at the same flow rate and for the same amount of time. All animals were monitored daily according to the PPL guidelines, two animals were excluded from the study following animal welfare guidelines. Echocardiography was performed 13 days post-surgery, followed by scarification and organ harvest on day 14 for further *ex vivo* analysis.

### *Ex vivo* measurements

Mice were sacrificed by cervical dislocation; the total body weight was recorded. Hearts were dissected using sterile surgical tools. Whole hearts were washed in PBS and weighed. Ventricular tissue was snap frozen in liquid nitrogen and stored at - 80 °C for future analysis.

To determine the tibia length, whole legs were dissected and digested in 0.8 M KOH overnight. Following the digestion, the tibia was measured and heart weight to tibia length ratio calculated.

### Immunofluorescence

Snap frozen cardiac tissue was embedded in O.C.T compound mounting medium (VWR Chemicals) for cryotome processing (10 μm thick cryosections). Samples were rinsed in 1x PBS and permeabilized with 0.2% Triton X-100 in PBS for 5 min. After another rinse in 1x PBS, the cardiac samples were blocked in 10% normal goat serum in PBS for 1h at room temperature. Samples were incubated in primary antibody (diluted in 1% BSA Gold Buffer: 20 mM Tris-HCl pH 7.5; 155 mM NaCl; 2 mM EGTA; 2 mM MgCl_2_) in a humidity chamber at 4°C overnight. Elisabeth Ehler (King’s College London) kindly gifted the anti-MYOM1 (1:50) antibody (clone B4, raised in mouse) for localisation of myomesin. For α-actinin, anti-α-actinin2 antibody (1:50, EP2529Y, abcam, rabbit) was used. Samples were rinsed in PBT (0.002% Triton X-100 in PBS) and washed three times for 5 min each in PBT, followed by secondary antibody incubation in a humidity chamber for 1 h at room temperature. For secondary antibody conjugates, anti-mouse Alexa Fluor 488 (1:100, A11017, life technologies) and anti-rabbit Alexa Fluor 568 (1:100, A21144, Life technologies) were used. After incubation, the samples were rinsed and washed three times in PBT for 5 min. Nuclei were counterstained using NucBlue (R37605, Invitrogen) according to the manufacturer’s guidelines. Coverslips were mounted with mounting media. Samples were analysed with Leica DM 6000 CFS Fluorescent Microscope using a 63x HCX PL AP0 oil objective.

Isolated cardiomyocytes were spun onto poly-lysine coated slides, fixed with 4% paraformaldehyde in PBS for 10 min and processed on the slides, apart from incubation of both primary and secondary antibodies overnight. Images were recorded on a Zeiss LSM880 Confocal Microscope with X63 Plan-APOCHROMAT water immersion lens at room temperature, using Zen 2.3 software.

Images were processed and analysed with ImageJ software.

### Histology

Paraffin-embedded hearts were cut into 7 μm thick sections. Samples were incubated in Histo- Clear and washed in 100% ethanol, followed by 95% ethanol for a few minutes. After rinsing in 1x PBS, sections were stained with hematoxylin for 30 s and rinsed under tap water before being counterstained with eosin for 30 s. After rinsing under tap water and in 1x PBS, samples were incubated in 95% ethanol and 100% ethanol. Before coverslips were applied, the samples were cleared of paraffin residues with Xylene. Images were taken using Axio Scan .Z1 slide scanner at 4x and 20x magnification in bright field mode using a 3CCD colour 2MP Hitachi 1200x1600 HV F202SCL camera.

### Contractility and Ca^2+^ measurements

Mouse left ventricular cardiomyocyte isolation was performed by Langendorff perfusion with Tyrode buffer (130 mM NaCl, 5.6 mM KCl, 3.5mM MgCl_2_), 5 mM 4-(2-hydroxyethyl)-1- piperazineethanesulphonic acid, HEPES, 0.4 mM Na_2_HPO_4_) containing 27 μg/ml liberase as previously described ^17^. Digestion of extracellular matrix by protease was quenched by diluting the resultant cell suspension with 3 volumes of 1% bovine serum albumin (BSA) in Tyrode solution and made sequentially Ca^2+^ competent by settling cells under gravity to form a loose pellet, removing the supernatant solution and adding back Tyrode with 1% BSA and 500 µM CaCl_2_, then 1% BSA containing 1 mM CaCl_2_ and finally resuspended in storage buffer (120 mM NaCl, 5.6 mM KCl, 5 mM MgSO_4_, 5 mM sodium pyruvate, 10 mM HEPES, 200 mM glucose, 200 mM taurine, 0.5 mM CaCl_2_) for further use.

Cell suspensions were divided, 10% was used for immunofluorescence slide preparation, 25% used for western blot sample preparation, 15% reserved for unloaded shortening measurements (without Ca^2+^ indicator loading) and the remaining 50% were incubated with fura2-AM-ester (fura2) for Ca^2+^ transient and pairwise unloaded sarcomere shortening measurements using IonOptix µstep apparatus. Cardiomyocyte contraction was assessed by fast Fourier transform of sarcomeric striations using phase-contrast microscopy. Ratiometric measurement of intracellular calcium ([Ca^2+^]_i_) transients were measured using dual excitation at 365 and 380 nm and measured emission at 520 nm by photomultiplier light acquisition.

Cardiomyocytes were loaded with fura2 (2 µM) in black Eppendorf tubes for 5 mins in Tyrode’s solution containing 250 µM CaCl_2_ + Pluronic F127, cells were washed for 10 mins in Tyrode’s containing 500 µM CaCl_2_ to remove any unloaded fura2 dye. At this point fura2 loaded cells were split with 50% of the cell suspension treated with 0.5 μM mavacamten and the remaining 50% treated with a DMSO vehicle control. Loaded cardiomyocytes were then placed in a glass-bottom chamber, mounted onto an inverted microscope, and perfused at 35±1 °C with 1.4 mM CaCl_2_ Tyrode’s solution containing 0.5 μM mavacamten or DMSO depending on prior treatment of the cells. Cardiomyocytes were field-stimulated at 40 volts and a frequency of 3 Hz for several minutes to allow steady-state [Ca^2+^]_i_ transients to be reached before recordings were taken. For each set of recordings, five background fluorescence measurements were taken from a cell-free field. [Ca^2+^]_i_ transients were measured as a function of the ratio of emissions from 365/380 excitation wavelength after subtraction of the background fluorescence (IonOptix) relative fluorescence was calibrated to [Ca^2+^] using the single exponential equation

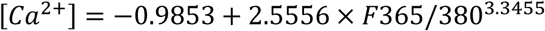

using the Ca^2+^ ionophore permeabilization method previously described ^18,19^. Average contractility and Ca^2+^ transients were determined from at least 10 individual raw tracings. Cells with poor morphology (e.g., excessive blebbing or asynchronous contraction) were not measured. Parameters were extracted using IonWizard 6.6.

### Mant-ATP assay and analyses

The Mant-ATP assay was performed on murine myocardial samples using a protocol adapted from Toepfer et al. ^20^.

In brief, after defrosting in permeabilization buffer (100 mM NaCl 8 mM MgCl_2_, 5 mM EGTA, 5 mM K_2_HPO_4_, 5 mM KH_2_PO_4_, 3 mM NaN_3_, 5 mM ATP, 1 mM DTT, 20 mM 2,3-butanedione monoxime, BDM, 0.1% Triton-X 100 at pH 7.0), the permeabilization of samples took place for 6 hrs on a rocker with exchange of the solution every 2 hrs. Samples were dissected following storage at −20°C in glycerinating solution (120 mM K acetate, 5 mM Mg acetate, 2.5 mM K_2_HPO_4_, 2.5 mM KH_2_PO_4_, 50 mM MOPS, 5 mM ATP, 20 mM BDM, 2 mM DTT, 50% glycerol (v/v), pH 6.8.) overnight. The myocardial tissue was then processed into ∼ 90 × 400 μm thin samples and placed in a chamber, followed by an additional permeabilization in permeabilization buffer for 10 min on ice and flushed using glycerinating buffer. The residual glycerol was replaced with rigor buffer (120 mM K acetate, 5 mM Mg acetate, 2.5 mM K_2_HPO_4_, 2.5 mM KH_2_PO_4_, 50 mM MOPS, 2 mM DTT at pH 6.8) for 10 min. Fluorescence acquisition to visualize fluorescent Mant-ATP wash-in was initiated by adding rigor buffer premixed with 250 μM Mant-ATP. Acquisition of the Mant-ATP chase was initiated after 10 min by the addition of ATP buffer (Rigor buffer + 4 mM ATP) to the chambers. A Nikon TE2000-E with a Nikon 20X/0.45 objective / Hammamatsu C9100 EM-CCD was used to visualize the fluorescence acquisition every 5 s / 20 ms over a time frame of 10 min in a DAPI filter setting. Mant-nucleotides were purchased from Invitrogen.

Fluorescence decay was measured in three independent areas per tissue sample. ImageJ ROI manager was used for the analysis. The subtraction of non-myosin bound Mant-ATP fluorescence signal was corrected by factor 52% as per Hooijman ^21^, the y-intercept describes the final fluorescence wash in data point. Data was normalized as initial fluorescent intensity and double exponential decay was calculated using the following equation:

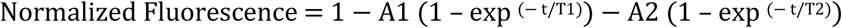

The initial rapid decay is represented by A1, and after non-specific Mant-ATP correction approximates the proportion of the disordered-relaxed state (DRX) of myosin heads, where T1 is the time constant. A2 indicates the slower decay resembling the super-relaxed state (SRX) of myosin heads and T2 is the time constant respectively.

### Western blotting

Western blots were performed using 4-15% mini-protean TGX gels (BioRad). For Phospho- Ser23/24 cTnI blots, a standard wet transfer on to PVDF membrane was used with subsequent blocking with 5% skimmed milk powder in TBST. Membranes were probed with a 1:2,000 dilution of anti Phospho-Ser23/24 TnI antibody (Cell Signalling) and 1:6,000 diluting of anti-rabbit-HRP secondary (Promega). Loading controls following antibody stripping with Restore Plus reagent (Thermo Fisher Scientific) used 1:5000 anti-cTnI (Abcam) with 1:15,000 anti-mouse-HRP (Promega) secondary and finally a 1:10,000 anti-GAPDH (Abcam) with a 1:30,000 anti-rabbit-HRP secondary. To measure RRXS/T protein kinase A target levels, a semi dry transfer was preformed using iBlot2 PVDF cassettes (Thermo Fisher Scientific), blocking and antibody incubation was performed in 5% BSA-TBST solution. A 1:1,000 dilution of anti-phospho-RRXS/T (Cell Signalling, ^22^) primary and 1:3,000 anti-rabbit-HRP secondary was used to probe for target proteins. An equal loading of all samples was run in parallel on a separate gel and stained with 5% Coomassie brilliant blue (Merck) to assess total protein loading. Densitometric measurements for all blots was performed using Image Lab following development of blot membranes with Dura ECL reagent (Thermo) using a GelDoc imaging system installed with software version 5.1 (BioRad). For Phospho-RRXS/T blots the entire lane was analysed using a band detection threshold of 95% and a cumulative densitometry was calculated. We excluded the over-exposed cTnI band at 24 kDa by manual removal. Coomassie stained loading controls showed a consistent actin band at 42 kDa which was analysed by densitometry as a loading control.

### Statistics

Data was analysed blind (to genotype and/or treatment groups). All values are given as mean ± standard error of mean (SEM). Data was tested for normality using the Kolmogorov-Smirnov test. To compare two unpaired sample groups, normally distributed data were analysed by Student’s t-test, and data that was not normally distributed were analysed by Mann-Whitney U-test. For comparison of three groups with normally distributed data, 1-way ANOVA followed by Tukey’s post-hoc test was used. For non-normally distributed data, Kruskal-Wallis test with Dunn’s test for multiple comparisons was used. Chronic adrenergic challenge response was analysed by 2-way ANOVA (with Tukey post-hoc test for multiple comparisons). For contractility and calcium transient measurements, all extracted parameters for each genotype and treatment group were split by individual mouse for hierarchical cluster statistical analysis using nested 1-way ANOVA as previously described ^23^.

All statistical analyses were performed blinded to genotype and treatment (where applicable) with GraphPad Prism 9.4.1.

Annotations used: * p < 0.05, ** p < 0.01, *** p < 0.001, **** p < 0.0001 versus WT, otherwise considered not significant (p > 0.05); n indicates number of animals in each group.

## Results

### Heterozygous *ALPK3tv* are enriched in patients with autosomal dominant adult onset HCM

We interrogated *ALPK3* genetic variants in a cohort of 230 HCM patients, in whom genetic screening had not identified a pathogenic genetic variant in sarcomere genes, and compared them to 6,219 controls without HCM. *ALPK3tv* were detected in 5 of 230 HCM cases (2.2%) and 9 of 6,219 controls (0.14%) (Fig. S1A, Table S1). This was a statistically significant excess of rare *ALPK3tv* in HCM cases compared to controls (OR 16.07, 95% CI 4.27-52.05; FET p= 0.0001), (Table S2). In contrast, there was no significant difference in the proportion of rare missense, non-truncating, or synonymous variants in HCM cases compared to controls (Table S2). This supports findings from recent studies ^8,9,13^ that *ALPK3tv* cause autosomal dominant adult onset HCM.

### A novel mouse model for *ALPK3tv*

One of the *ALPK3tv* from our HCM cohort (Figure S1A, Table S1) was used to generate a mouse model for *ALPK3*-associated HCM, *Alpk3* K201X (Figure S1B, C). We chose a human autosomal dominant *ALPK3tv*, as opposed to a full gene knockout ^24^, to account for potential dominant negative effects of the truncated protein, which in case of truncation at Lysine 201 would lack the putative kinase domain. Since recessive (e.g. homozygous) *ALPK3tv* can lead to early onset cardiomyopathy in patients, the cardiac structure and function of homozygous (Hom) mice with the *Alpk3* K201X truncation variant were analysed and compared to wildtype (WT) animals alongside heterozygous (Het) animals using echocardiography (Figs. 1, S2, Table S3).

Homozygous *Alpk3* K201X animals were viable and fertile, and showed no gross extra- cardiac abnormalities. Their hearts displayed a significant impairment of systolic function compared to WT mice at 3 months, evidenced by a significant reduction of fractional shortening (Fig. 1A) and peak systolic velocity (Fig. 1B). Moreover, these mice displayed features of hypertrophy, e.g. increased left ventricular mass (Fig. 1C) on echocardiography. In support, homozygous mouse hearts were enlarged (Fig. S2A) and had increased heart weight to tibia length ratio on gravimetry (Figs. 1D), but wall thicknesses were preserved (Table S3). Ventricular dilatation was evidenced by increased internal diameter of the left ventricle in both diastole and systole (Fig. 1E, F). These results suggest that the newly generated mouse model recapitulates expected cardiac features of human recessive *ALPK3tv*, namely cardiac hypertrophy and systolic dysfunction.

At molecular level, the *Alpk3* K201X variant undergoes partial nonsense-mediated decay, with only ∼27 % of *Alpk3* transcript level in homozygous *Alpk3* K201X hearts compared to WT (Fig. S1C). Moreover, homozygous hearts displayed induction of transcripts related to the foetal gene programme and hypertrophic signalling in support of cardiomyopathy in these hearts (Fig. S3).

A side-by-side comparison of our novel mouse with the previously published *Alpk3* knockout (KO) model ^2^ showed identical phenotypes on echocardiography (Table S4) and qPCR (Fig. S3).

### Heterozygous mice have normal baseline cardiac structure and function

Mice heterozygous for *Alpk3* K201X demonstrated normal cardiac function with no signs of hypertrophy or dilatation at 3 months. The analysed hearts were indistinguishable from WT littermates (Figs. 1, S2, Table S3).

In agreement with the absence of a phenotype on echocardiography, mice heterozygous for *Alpk3* K201X showed no induction of the foetal gene programme or transcripts related to hypertrophy on qPCR (Fig. S3). Histology was unremarkable in both heterozygous and homozygous setting (Fig. S4A), and there was no molecular evidence of fibrosis (Fig. S4B). However, isolated cardiomyocytes from both heterozygous and homozygous *Alpk3* K201X mice showed an increase in cell length, while cell width was normal (Fig. S4C).

As a role for ALPK3 in M-band organisation was implicated previously ^2^, we analysed localisation of the M-band protein myomesin using immunofluorescence staining on cardiac tissue and isolated cells, but failed to detect any differences between the genotypes (Fig. S5). Please note that there is currently no working antibody available to localise Alpk3.

As HCM often manifests in adulthood in patients, adult mice of 6 months age were also investigated. Animals showed normal cardiac structure and function on echocardiography also at this age (Fig. 1, Table S3). Moreover, heart weight relative to tibia length was normal on gravimetry (Table S3).

In summary, mice heterozygous for *Alpk3* K201X failed to show signs of cardiomyopathy.

### Chronic adrenergic challenge aggravates the hypertrophic response in heterozygous mice

Since heterozygous *Alpk3tv* mice did not display an overt phenotype, we subjected animals to a chronic adrenergic challenge to induce a hypertrophic response via isoprenaline and phenylephrine (Iso/PE) treatment. The hypertrophic response of the murine hearts was examined upon two weeks of adrenergic stimulation using echocardiography. The heart weight to tibia length ratio increased in both WT and heterozygous animals upon drug-treatment compared to their control groups, confirming the treatment induced cardiac hypertrophy (Fig. 2A). There was a larger response in heterozygous *Alpk3* K201X hearts compared to WT hearts. In agreement, the calculated left ventricular mass based on echocardiography was enlarged in drug-treated heterozygous animals compared to corresponding control group and drug-treated WT animals (Fig. 2B). Left ventricular anterior wall thickness was increased in drug-treated heterozygous animals compared to control animals of the same genotype (Fig. 2C). However, systolic function was unchanged in both genotypes upon drug-treatment (ejection fraction and fractional shortening in Tab. S5). This experiment indicates that the mice carrying the heterozygous *Alpk3tv* have an aggravated hypertrophic response to chronic adrenergic challenge, while their systolic function is preserved.

**Figure 2.**
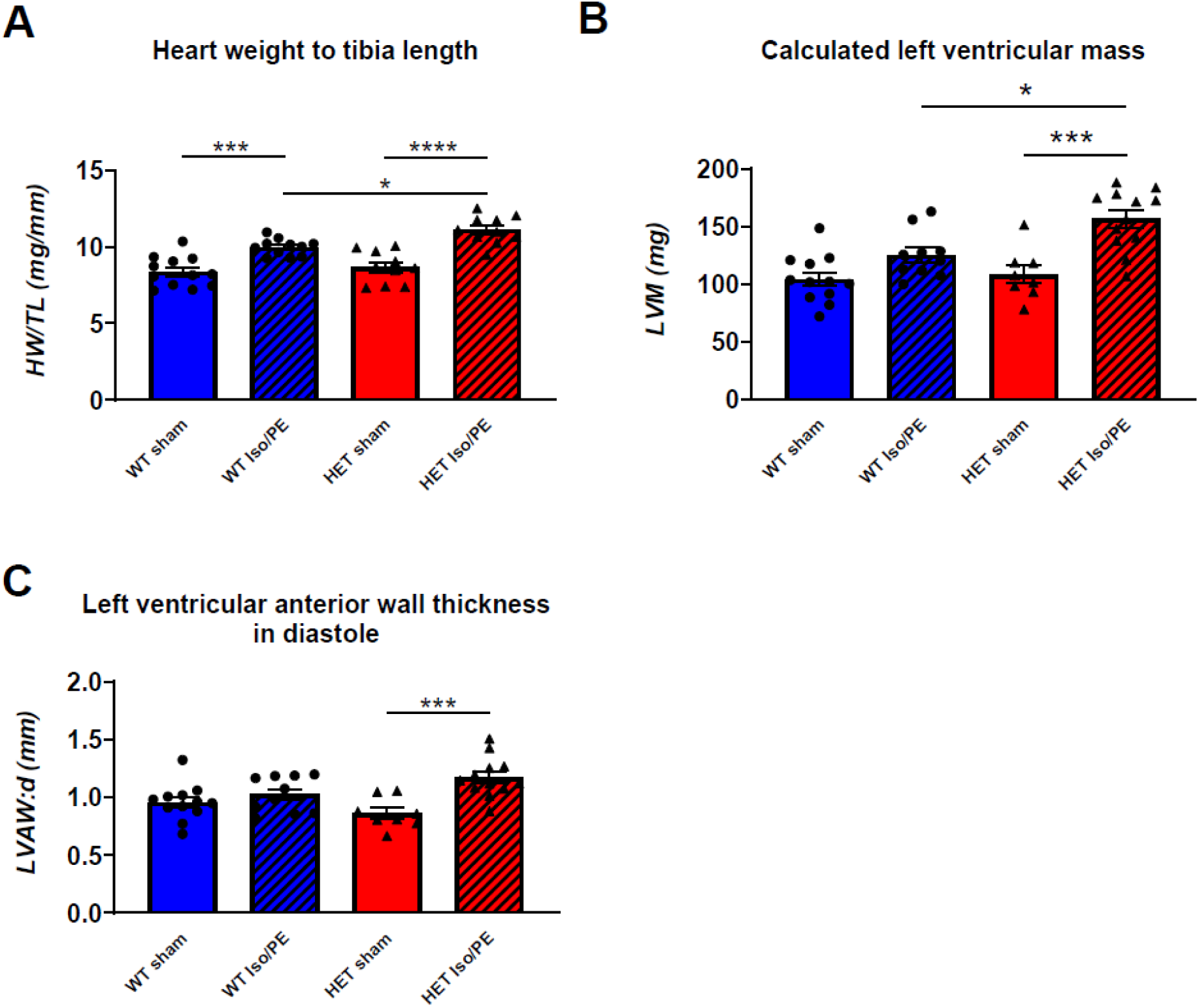
Chronic adrenergic challenge aggravates the hypertrophic response in heterozygous *Alpk3 K201X*. (**A**) The heart weight (HW) to tibia length (TL) ratio increased in both wildtype (WT) and heterozygous (Het) animals upon isoprenaline and phenylephrine (Iso/PE) treatment compared to their control groups (sham) treated with saline, indicating cardiac hypertrophy. There was a stronger response in Iso/PE treated Het compared to WT animals. WT sham n=12; WT Iso/PE n=11; Het sham n=10; Het Iso/PE n=10. (**B**) Het *Alpk3* K201X mice displayed an aggravated hypertrophic response upon chronic adrenergic challenge as evidenced by calculated left ventricular mass (LVM). (**C**) Left ventricular anterior wall thickness in diastole (LVAW:d) increased in challenged Het animals only. WT sham n=12; WT Iso/PE n=10; Het sham n=8; Het Iso/PE n=12 (for B and C). Values are presented as mean ± SEM. * p < 0.05, ** p < 0.01, **** p < 0.0001 (2-way-ANOVA), otherwise considered not significant. For a full set of echocardiographic parameters please refer to Table S5.

### Cardiomyocytes from *Alpk3* K201X mice are hypercontracted with prolonged relaxation and reduced fractional shortening

Sarcomeric HCM is associated with altered cardiomyocyte contractility and calcium transients. This is well documented for mouse models of HCM, even when subtle or no overt cardiac phenotypes are observed *in vivo* ^25,26^. In order to investigate a potential cellular phenotype in our model, we isolated adult left ventricular cardiomyocytes from heterozygous and homozygous mice and compared their unloaded contractile parameters to cells isolated from WT mice. In cells preloaded with fura2, we observed a significant and dose dependent decrease in basal sarcomere length of 4.3 ± 0.3 and 7.4 ± 0.3% and an increase in the time to 90% relaxation (T90_relaxation_) by 23.2 ± 3.2 and 62 ± 3 ms for heterozygous and homozygous cells respectively compared to WT (Figs. 3A, C, F, S7, Table S6A). Homozygous cells also showed a 24.6 ± 1.3 ms prolongation in contraction (Figs. 3A, E, S7, Table S6A). These data are indicators of hypercontractility and diastolic dysfunction. These changes were observed in both the presence and absence of calcium indicator fura2 (Figure S6, Table S6C). We also observed a 22 ± 3% reduction in contractile magnitude (as measured by fractional shortening) in heterozygous cells but no change in homozygous cells in the presence of fura2 (Figs. 3A, D, S7, Table S6A). However, in dye free measurements, there was no change in heterozygous cells and a 36 ± 5% drop in contractile magnitude for homozygous cells (Fig. S6A, C, Table S5C). The discordance in the data can be attributed to contractile impairment caused by fura2 and the observation made in dye free measurements gives confidence that systolic dysfunction evidenced by reduced fractional shortening is a real effect.

**Figure 3.**
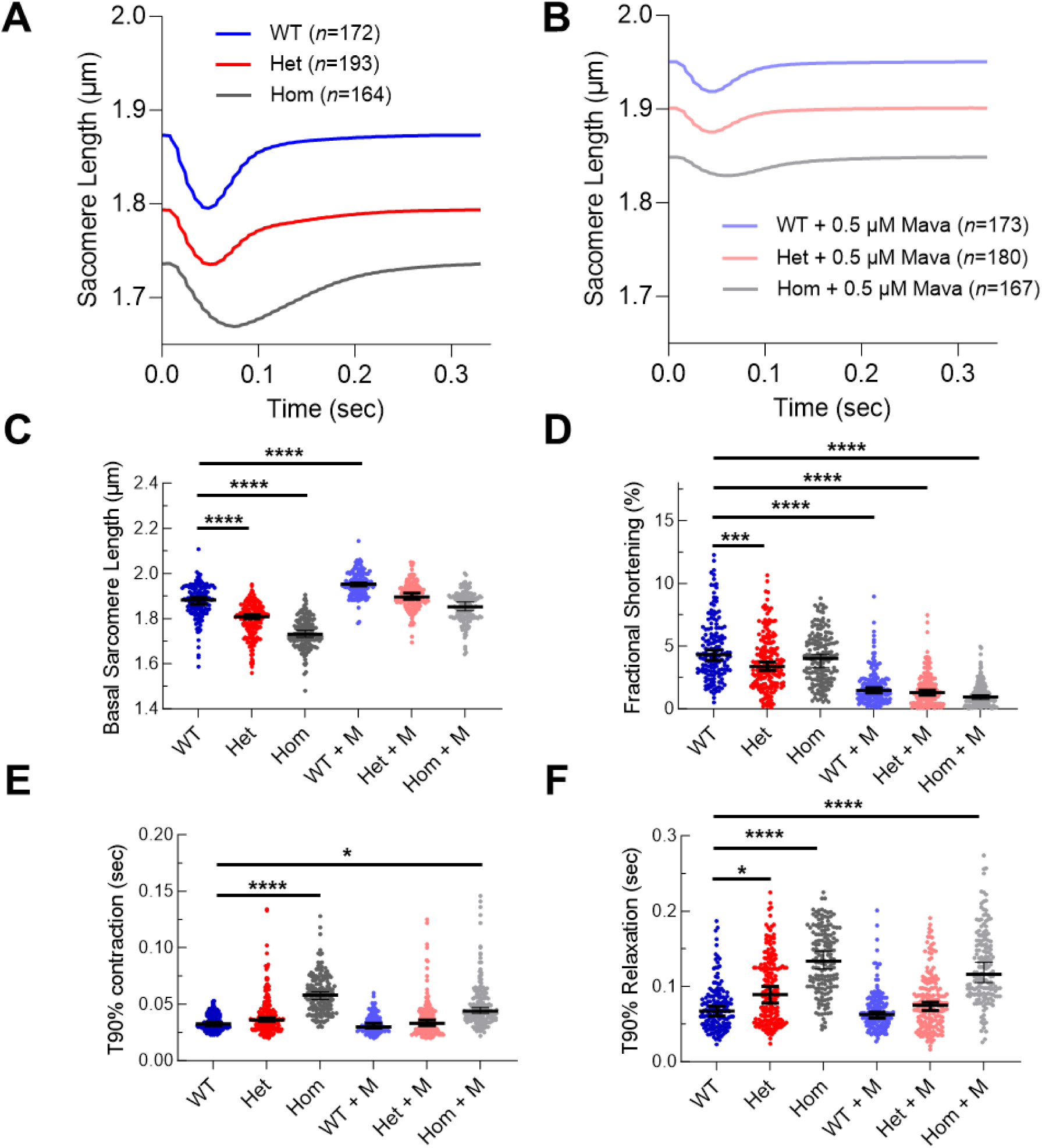
Isolated left ventricular cardiomyocytes from *Alpk3* K201X mice display hypercontractile unloaded sarcomere shortening with prolonged relaxation, which is partially rescued by mavacamten treatment. (**A**) Average sarcomere length traces from heterozygous *Alpk3* K201X (Het, red) and homozygous *Alpk3* K201X cardiomyocytes (Hom, grey) decreased compared to wild type (WT) cardiomyocytes (blue). (**B**) Sarcomere length traces following 0.5 μM mavacamten treatment revealed an increase in sarcomere length across all genotypes. Dot plots for selected extracted parameters: basal sarcomere length (**C**), fractional shortening (**D**), time to 90 % (T90%) contraction (**E**) and T90% relaxation (**F**) demonstrated partial rescue of contractile parameters in Het and Hom cardiomyocytes. Black lines represent the median ± 95% confidence interval. Statistical differences comparing WT to all other groups were calculated using nested 1-way ANOVA to adjust for hierarchical clustering of individual data sets between mice (Figure S6). *p<0.05, ***p<0.001, ****p<0.0001. Sample size n (cells per animal group) stated in the figure.

### Reduced protein kinase A dependent protein phosphorylation as a potential mechanism for contractile defects in *Alpk3* K201X cardiomyocytes

In order to examine how a truncation in ALPK3, a protein with no known sarcomeric role, may be mediating an effect on contractility, we investigated the phosphorylation status of cardiac troponin I (cTnI). We found chronic reduction in serine23/24 cTnI modification (Fig. 4A,B), which is known to slow relaxation and sensitise the myofilament ^27^. As this is a known site for phosphorylation by protein kinase A (PKA) ^28^, we also assessed global PKA substrate status. We observed a dose dependent reduction in all potential PKA target proteins (Fig. 4C). Cumulative densitometric analysis (Fig. 4D) and normalisation to a cardiac actin loading control showed that the level of PKA dependent phosphorylation is reduced by 39.3 ± 3.7% and 50.1 ± 2.6% in heterozygous and homozygous cells respectively (Fig. 4E).

**Figure 4.**
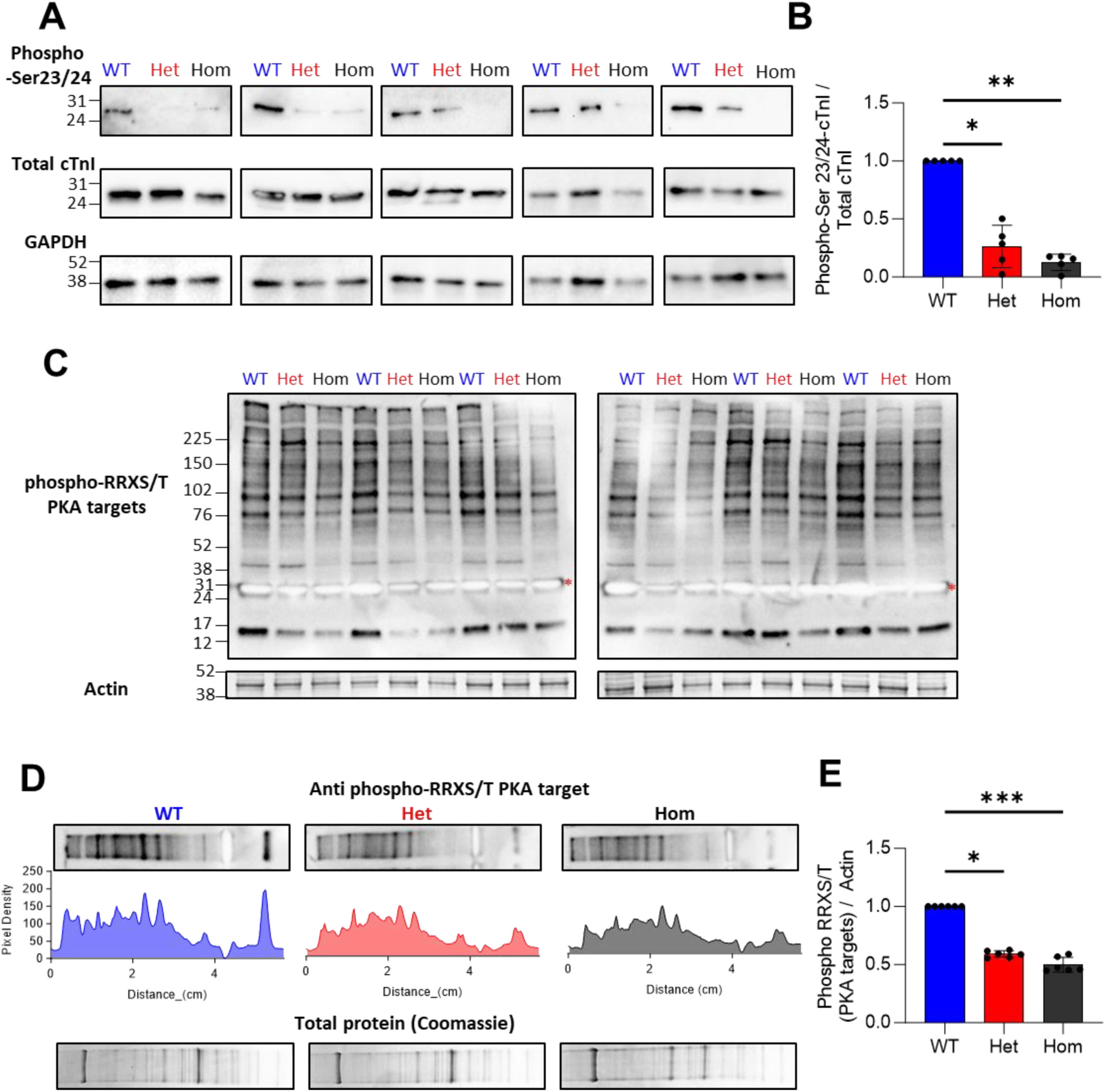
Depleted PKA dependent protein phosphorylation in *Alpk3* K201X mice. Western blots of isolated cardiomyocyte preparations showed the phosphorylation of cTnI at Ser23/24 compared to total cTnI and GAPDH loading control (**A**), Densitometry analysis in (**B**) showed the degree of phosphorylation in wild type (WT), heterozygous (Het) and homozygous (Hom) mice (n=5 mice per group). All PKA dependent phosphorylation targets were detected using an anti-phospho-RRXS/T antibody (**C** top), red stars denote an overexposed phospho-cTnI band, which was manually removed from the subsequent densitometric analysis. The loading control was cardiac actin, taken from the 42 kDa band on Coomassie stained control gels. (**D**) shows examples of lane densitometry measurements used to calculate cumulative band intensity across all detected proteins for WT, Het and Hom mice. The ratio of PKA target densitometry versus cardiac actin from Coomassie stained gels run in parallel is plotted in (**E**) (n=6 mice per group). * p<0.05, ** p<0.01, *** p<0.001 using a Kruskal-Wallis test with Dunn’s test for multiple comparisons.

### Abnormal calcium transients highlight complex changes to excitation-contraction coupling in cardiomyocytes from *Alpk3* K201X mice

Calcium transients from fura2 loaded cardiomyocytes showed dose-dependent increases in diastolic intracellular calcium by ∼50 % for heterozygous and ∼65 % for homozygous mice compared to WT (Figs. 5A, C S7, Table S6B). These changes are concordant with baseline contractile changes. Homozygous cells also showed a 2.3 ± 0.8 ms prolongation in time to 90% release (T90_release_) and an increase of 22.0 ± 1.5 ms in time to 50% reuptake (T50_reuptake_) (Figs. 5A, E, F, S7, Table S6B) which parallels changes in the contraction and relaxation velocity. However, the calcium transient amplitude remained unchanged across all genotypes (Figs. 5A, D, S7, Table S6B) in contrast to observations of reduced fractional shortening. This highlights a paradoxical desensitisation of the contractile apparatus to calcium on the background of hypercontractility, which may be unique to *ALPK3* HCM.

**Figure 5.**
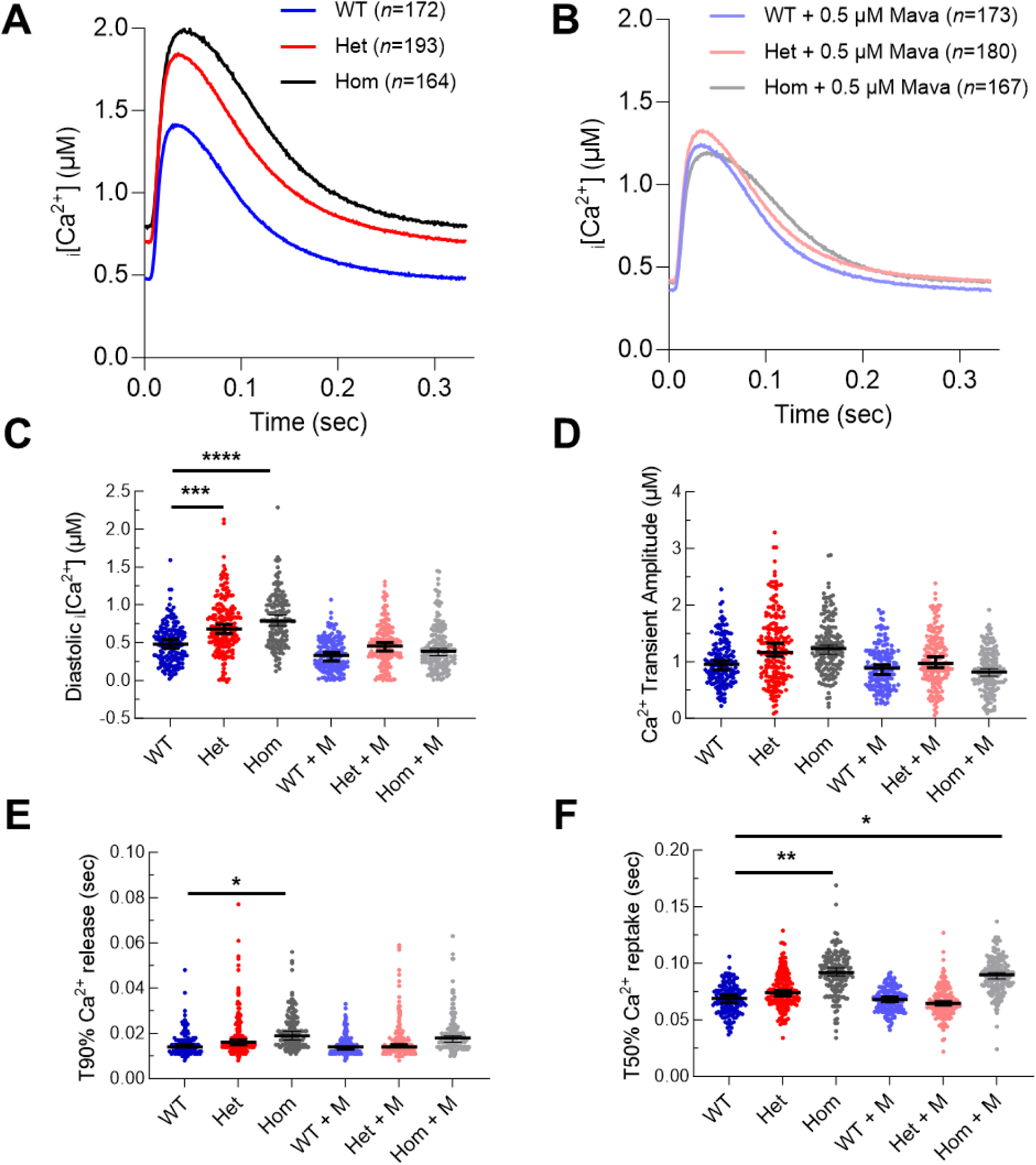
Isolated left ventricular cardiomyocytes from *Alpk3* K201X mice have elevated diastolic intracellular calcium levels and prolonged calcium reuptake, which is fully rescued by mavacamten treatment. (**A**) Average calibrated calcium transients from wild type (WT, blue), heterozygous *Alpk3* K201X (Het, red), homozygous *Alpk3* K201X cardiomyocytes (Hom, grey). (**B**) Calcium transients following treatment with 0.5 μM mavacamten were similar between all genotypes. Dot plots for selected extracted parameters: Diastolic intracellular calcium levels [Ca^2+^]_i_ (**C**), calcium transient amplitude (**D**), time to 90 % (T90%) calcium release (**E**) and time to 50 % (T50%) calcium reuptake (**F**). Black lines represent median ± 95% confidence interval. Statistical differences comparing WT to all other groups were calculated using nested 1-way ANOVA to adjust for hierarchical clustering of individual data sets between mice (Figure S6I-P). *p<0.05, **p<0.01, ***p<0.001, ****p<0.0001. Individual n numbers (cells per group of animals) and all extracted parameters are tabulated in supplementary Table S6C.

### Myosin conformation changes in *Alpk3* K201X mice are consistent with hypercontractility and HCM

A reduction in the ratio of super-relaxed state (SRX) to disordered-relaxed state (DRX) of myosin heads is a molecular hallmark of sarcomeric HCM, observed in cardiac tissue from human patients and in animal models ^29^. To probe whether mice with the *Alpk3* K201X variant have changes in the states of myosin heads, myosin ATP binding was investigated using cardiac tissue from WT, heterozygous and homozygous mice.

Heterozygous mice showed a significantly reduced proportion of myosin heads in the SRX state, compared to WT littermates (Fig. 6). This suggests molecular changes occurring in the heterozygous *Alpk3* K201X mice are consistent with HCM and are more pronounced in homozygous (Hom) animals, indicating a dose-dependent effect.

**Figure 6.**
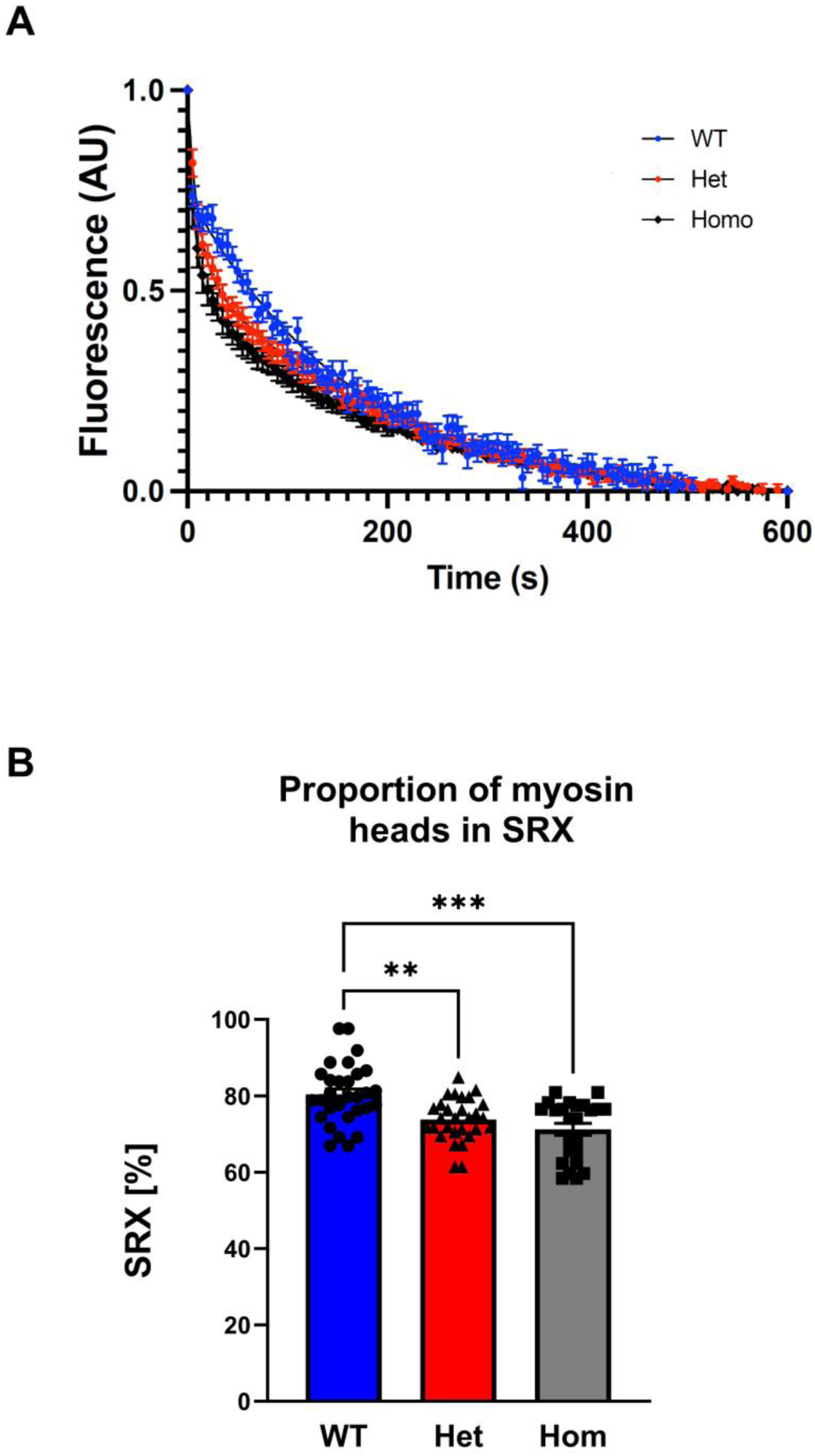
Molecular changes in the heterozygous mice are consistent with HCM. Myosin ATP binding was analysed via a Mant-ATP assay on cardiac tissue of 3-month-old *Alpk3 K201X* mice. (**A**) – Averaged curves of Mant-ATP hydrolysis for cardiac tissue from wildtype (WT), heterozygous (Het) and homozygous (Hom) *Alpk3 K201X* mice. (**B**) Despite the absence of cardiac functional or structural changes on echocardiography Het *Alpk3 K201X* mice showed a significantly reduced proportion of myosin heads in the super-relaxed state (SRX), compared to WT littermates. This was observed to be even more significantly reduced in Hom animals. Values are presented as mean ± SEM. ** p < 0.01, *** p < 0.001 versus WT (Kruskal-Wallis test, n=4-6 technical replicates, n=3 biological replicates per genotype).

### Mavacamten partially improves the cellular phenotype

Mavacamten is an allosteric myosin ATPase inhibitor that has been shown to reduce DRX myosin, thereby reversing pathological changes in HCM ^20^. Given the observed reduction of the SRX state of myosin heads in our mice, we treated isolated cardiomyocytes from our mouse model with 0.5 µM mavacamten and repeated contractility and calcium transient measurements to probe for a beneficial effect of mavacamten. Treatment of WT cells showed increased basal sarcomere length and reduced fractional shortening (Fig. 3A-D) with unchanged diastolic calcium levels and transient amplitude (Fig. 5A-D) as previously described ^30,31^. When treating both heterozygous and homozygous *Alpk3* K201X cells, mavacamten rescued the calcium transients to more closely resemble WT flux by reducing diastolic Ca^2+^ levels and Ca^2+^ reuptake time. The exception to this was for calcium reuptake speed in treated homozygous mice cells where T50_reuptake_ was prolonged by 20.5 ± 1.2 ms compared to WT (Figs. 5B, F, S7, Table S6B). Mavacamten treatment also rescued basal sarcomere length (Figs. 3C, S6C, S7, Table S6A, C). Because *Alpk3* K201X cells have impaired fractional shortening, mavacamten treatment exacerbates contractile impairment by 67.6 ± 1.9% and 75.5 ± 1.5% for heterozygous and homozygous cells, respectively, compared to 62 ± 2% reduction in mavacamten treated WT cells (Fig. 3D). Contraction and relaxation velocity were still prolonged in treated homozygous cells by 13.6 ± 1.8 and 48 ± 4 ms, respectively, compared to WT (Fig. 3E, F, S7, Table S6A) indicating that the genotype effect still predominates. The partial rescue of *Alpk3* variant cells’ contractile parameters by mavacamten were recapitulated in the absence of fura2 loading in qualitatively the same manner (Figure S6, Table S6C).

In summary, mavacamten can partially restore abnormal calcium handling in the mouse model and has beneficial effects on resting sarcomere length and relaxation.

## Discussion

Using genetic screening, we confirmed that heterozygous *ALPK3tv* can cause autosomal dominant HCM. Case-control analysis revealed an excess of rare *ALPK3tv* in our HCM case cohort, compared to controls. The level of enrichment observed in our cohort (2.0%; OR 16.07, 95% CI 4.27-52.05) was comparable to that reported in recent studies (1.56%; OR 16.17, 95% CI 10.31-24.87) ^9^. This indicates that while heterozygous *ALPK3tv* are causal for HCM, they have a lower odds ratio, reflecting lower penetrance, than seen with typical sarcomeric HCM genes. This is in keeping with the lack of overt HCM in the majority of the heterozygous carriers in the *ALPK3tv* families first described with recessive cardiomyopathy.

In this study we have generated a novel mouse model incorporating the *Alpk3* K201X variant based on a patient’s rare heterozygous *ALPK3tv* identified in our HCM cohort. This variant was chosen for its amino-terminal position, expected to produce (if at all) a truncated ALPK3 protein without the (pseudo-)kinase domain. Reduced transcript levels for *Alpk3* were observed in a dose dependent manner, reaching significant reduction in homozygous mice (Figure S1C). This argues for at least partial haplo-insufficiency through nonsense-mediated decay. In the absence of a working antibody for ALPK3, the consequences on the Alpk3 protein, e.g. presence of truncated protein, remain unknown.

In the homozygous setting, the mouse model recapitulates observations of clinical reports of biallelic carriers of *ALPK3tv*, leading to severe, often lethal, paediatric onset cardiomyopathy ^8,11,12^: The homozygous *Alpk3* K201X mice have impaired systolic function and hypertrophy; we found their phenotype indistinguishable from *Alpk3* knockout mice. In our hands, both models survive well into adulthood. This is in contrast to a recent report on the same *Alpk3* knockout mice ^2^, which – despite similar cardiac observations of systolic dysfunction and hypertrophy – did not survive beyond 14 weeks. We can only speculate whether genetic background (C57bl/6J versus C57bl/6N) or environmental factors, e.g. diet, health status and microbiome ^32^, could be responsible for the observed differences between laboratories.

Despite our aim to mirror the pathological human *ALPK3tv* found in our patient cohort, the heterozygous mice showed no overt cardiac phenotype at 3 or 6 months of age. This agrees with a recent report using an *Alpk3* null mouse model: mice with a heterozygous deletion of *Alpk3* developed hypertrophy only when aged beyond 1 year of age ^2^. In addition, McNamara *et al.* reported normal cardiac dimensions and function in 3-week-old mice heterozygous for the c.5294G>A, p.W1765X *ALPK3* patient-specific variant, whereas mice homozygous for this variant presented with pronounced systolic and diastolic dysfunction as well as left ventricular hypertrophy ^4^.

Moreover, due to differences in cardiac physiology and lack of stressors, the full disease spectrum of humans often cannot be fully reflected in animal models. This has also been reported for other heterozygous cardiomyopathy gene models, such as *Csrp3, Mybpc3, Myh7* and *Ttn* displaying phenotypes in homozygous settings only, whilst heterozygous animals showed no cardiac abnormalities ^17,33–35^. Interestingly, individuals carrying heterozygous *ALPK3tv*, as family members of patients with recessive *ALPK3tv*, often have no clinical characteristics of HCM; only three out of 21 patients showed clinical signs of HCM ^6,8,12^. Nevertheless, heterozygous *Alpk3* K201X mice showed an aggravated hypertrophic response to chronic adrenergic stimulation. This mirrors the observations of a *Mypbc3* HCM mouse model, where the same Iso/PE treatment helped to unmask disease features in the heterozygous mice ^36^.

Despite the lack of baseline phenotype *in vivo*, isolated cardiomyocytes from both heterozygous and homozygous *Alpk3* K201X mice showed striking abnormalities: they displayed reduced diastolic sarcomere length, accompanied by an increase in diastolic calcium concentration. Moreover, they showed prolonged relaxation, backed up by delayed calcium re-uptake into the sarcoplasmic reticulum. These are hallmarks of cardiomyocytes from sarcomeric HCM models ^19,30^ and effects were dose-dependent, i.e. more pronounced in homozygous than in heterozygous *Alpk3* K201X cells. In contrast to classical sarcomeric HCM models, the *Alpk3* K201X cells displayed some indications of reduced fractional shortening. While this fits with the systolic dysfunction observed *in vivo* for the homozygous mice, reduced contractility at a cellular level is usually observed in models of DCM ^37^. Therefore, we demonstrate for the first time, that the *ALPK3tv* – as a non-sarcomeric HCM variant – causes a ‘mixed model’ of sensitization and desensitization of the myofilament, combining features of cardiomyocytes from both HCM and DCM models.

Moreover, we observed reduced PKA-mediated phosphorylation in *Alpk3* K201X cardiomyocytes, including markedly reduced phosphorylation of troponin I at serine23/24. The latter is predicted to result in increased troponin Ca^2+^ affinity and hence, at least in part, may be responsible for the observed slower relaxation and reduced sarcomere length ^27^. Moreover, reduced global PKA substrate phosphorylation may implicate changes to a range of other functional targets that may modulate contractility and Ca^2+^ handling ^38^. Whether these observed functional changes are caused by ALPK3 directly regulating the activity and/or localisation of PKA and/or phosphodiesterases to alter phosphorylation levels, or are part of a chronic adaptation of the myofilaments remains unknown. In support of the latter, a more acute phospho-proteome study using *ALPK3* mutant iPSC-derived cardiomyocytes cultured for a maximum of 30 days did not observe a reduction in phosphorylation of PKA-targets on the myofilament ^4^.

Reduced SRX state of myosin heads is an important molecular hallmark of HCM ^39^. Biochemically it can be assessed by the Mant-ATP assay, which has previously documented HCM-associated changes in myosin ATPase kinetics, i.e. a shift from SRX to DRX, for pathogenic *MYBPC3* and *MYH7* variants, in mouse models, cellular models, in-silico models and human tissue samples ^20,29,40–42^.

Although *Alpk3* K201X heterozygous mice lack an overt phenotype, they show a reduction in the biochemically defined SRX/DRX ratio by Mant-ATP; which became more pronounced in homozygous mice. The reduction of myosin SRX in the *Alpk3* K201X highlights a shared phenotype with other common HCM variants studied using mouse models. However, it is worth noting that the observed changes in the SRX/DRX ratio are less striking than in a *Mybpc3* mouse model ^20^.

Mavacamten is a myosin ATPase inhibitor, known to stabilise the SRX state ^43^. It is now used to treat symptomatic obstructive HCM in patients. Given the reduced SRX/DRX ratio in our mouse model and the observed HCM-like dysfunction at cellular level, we tested whether mavacamten could restore calcium handling and contractility in the cardiomyocytes. Mavacamten rescued the reduced diastolic sarcomere length and partially rescued the impaired relaxation of *Alpk3* K201X cardiomyocytes by alleviating changes in calcium handling. However, it had detrimental effects on fractional shortening, which parallels observations on HCM-causing variants in thin filament proteins, e.g. cardiac troponin T R92Q and cardiac troponin I R145G ^30^.

Therefore, while we demonstrate here for the first time that mavacamten may have beneficial effects on patients with *ALPK3tv*, careful dose titration and tight surveillance might be required to prevent excessive reductions in systolic function in those patients. Future *in vivo* animal studies and clinical trials are needed to evaluate the benefits and risks of mavacamten treatment in this patient group.

In conclusion, our novel *Alpk3* K201X mouse model for HCM caused by autosomal dominant *ALPK3tv* demonstrates hallmarks of HCM at the level of isolated cardiomyocytes. To our knowledge, this is the first illustration of a ‘non-sarcomeric’ HCM disease gene manifesting such effects. However, our findings indicate more complex molecular changes in the presence of *ALPK3tv*, i.e. a mixed picture of sensitization and desensitization of the myofilament, with features of both HCM and DCM seen, in keeping with the clinical phenotype. We further demonstrate that mavacamten can restore some aspects of cellular dysfunction and may therefore be a therapeutic option for patients with *ALPK3tv*.

## Supporting information

Electronic Supporting Information

## Declarations

### Funding

This work was funded by the British Heart Foundation (PG/19/45/34419, to LL, KG, CR and HW), by The Medical Research Council (MR/V009540/1, to SBS and KG), by the National Centre for the 3Rs and Industry (NC/T001747/1 to KG), by a NIHR and Health Education England (HEE) Healthcare Science Doctoral Research Fellowship (NIHR-HCS-D13-04-006 to KT), by a Sir Henry Dale Wellcome Fellowship (222567/Z/21/Z to CNT), by a BHF Centre of Research Excellence Intermediate Transition Fellowship (RE/18/3/34214 to CNT) and by the NIHR Oxford Biomedical Research Centre (to EO). The Institute of Cardiovascular Sciences, University of Birmingham, has received an Accelerator Award by the British Heart Foundation (AA/18/2/34218). CS is the recipient of a National Health and Medical Research Council (NHMRC) Investigator Grant (#2016822) and a New South Wales Health Cardiovascular Disease Clinician Scientist Grant. RDB is the recipient of a New South Wales Health Cardiovascular Disease Senior Scientist Grant.

### Conflicts of interest

On behalf of all authors, the corresponding author states that there is no conflict of interest.

### Availability of data and material

Data underlying this article will be shared on reasonable request to the corresponding author.

## Notes

### Competing Interest Statement

The authors have declared no competing interest.

## References

1 Middelbeek, J., Clark, K., Venselaar, H., Huynen, M. A. & van Leeuwen, F. N. The alpha-kinase family: an exceptional branch on the protein kinase tree. Cell Mol Life Sci 67, 875–890, doi:10.1007/s00018-009-0215-z (2010).

2 Agarwal, R. et al. Pathogenesis of Cardiomyopathy Caused by Variants in ALPK3, an Essential Pseudokinase in the Cardiomyocyte Nucleus and Sarcomere. Circulation, doi:10.1161/CIRCULATIONAHA.122.059688 (2022).

3 Feng, W., Bogomolovas, J., Wang, L., Li, M. & Chen, J. ALPK3 Functions as a Pseudokinase. Circulation 148, 1911–1913, doi:10.1161/CIRCULATIONAHA.123.065993 (2023).

4 McNamara, J. W. et al. Alpha kinase 3 signaling at the M-band maintains sarcomere integrity and proteostasis in striated muscle. Nature Cardiovascular Research 2, 159–173, doi:10.1038/s44161-023-00219-9 (2023).

5 Hosoda, T. et al. A novel myocyte-specific gene Midori promotes the differentiation of P19CL6 cells into cardiomyocytes. J Biol Chem 276, 35978–35989, doi:10.1074/jbc.M100485200 (2001).

6 Almomani, R. et al. Biallelic Truncating Mutations in ALPK3 Cause Severe Pediatric Cardiomyopathy. J Am Coll Cardiol 67, 515–525, doi:10.1016/j.jacc.2015.10.093 (2016).

7 Aung, N. et al. Genome-Wide Analysis of Left Ventricular Image-Derived Phenotypes Identifies Fourteen Loci Associated With Cardiac Morphogenesis and Heart Failure Development. Circulation 140, 1318–1330, doi:10.1161/CIRCULATIONAHA.119.041161 (2019).

8 Herkert, J. C. et al. Expanding the clinical and genetic spectrum of ALPK3 variants: Phenotypes identified in pediatric cardiomyopathy patients and adults with heterozygous variants. Am Heart J 225, 108–119, doi:10.1016/j.ahj.2020.03.023 (2020).

9 Lopes, L. R. et al. Alpha-protein kinase 3 (ALPK3) truncating variants are a cause of autosomal dominant hypertrophic cardiomyopathy. Eur Heart J 42, 3063–3073, doi:10.1093/eurheartj/ehab424 (2021).

10 Phelan, D. G. et al. ALPK3-deficient cardiomyocytes generated from patient-derived induced pluripotent stem cells and mutant human embryonic stem cells display abnormal calcium handling and establish that ALPK3 deficiency underlies familial cardiomyopathy. Eur Heart J 37, 2586–2590, doi:10.1093/eurheartj/ehw160 (2016).

11 Caglayan, A. O. et al. ALPK3 gene mutation in a patient with congenital cardiomyopathy and dysmorphic features. Cold Spring Harb Mol Case Stud 3, doi:10.1101/mcs.a001859 (2017).

12 Jaouadi, H. et al. Novel ALPK3 mutation in a Tunisian patient with pediatric cardiomyopathy and facio-thoraco-skeletal features. J Hum Genet 63, 1077–1082, doi:10.1038/s10038-018-0492-1 (2018).

13 Dai, J. et al. The Involvement of ALPK3 in Hypertrophic Cardiomyopathy in East Asia. Front Med (Lausanne) 9, 915649, doi:10.3389/fmed.2022.915649 (2022).

14 Thomson, K. L. et al. Analysis of 51 proposed hypertrophic cardiomyopathy genes from genome sequencing data in sarcomere negative cases has negligible diagnostic yield. Genet Med 21, 1576–1584, doi:10.1038/s41436-018-0375-z (2019).

15 Richards, S. et al. Standards and guidelines for the interpretation of sequence variants: a joint consensus recommendation of the American College of Medical Genetics and Genomics and the Association for Molecular Pathology. Genet Med 17, 405–424, doi:10.1038/gim.2015.30 (2015).

16 Sigmon, J. S. et al. Content and Performance of the MiniMUGA Genotyping Array: A New Tool To Improve Rigor and Reproducibility in Mouse Research. Genetics 216, 905–930, doi:10.1534/genetics.120.303596 (2020).

17 Jiang, H. et al. Functional analysis of a gene-edited mouse model to gain insights into the disease mechanisms of a titin missense variant. Basic Res Cardiol 116, 14, doi:10.1007/s00395-021-00853-z (2021).

18 Carnicer, R. et al. Cardiomyocyte GTP cyclohydrolase 1 and tetrahydrobiopterin increase NOS1 activity and accelerate myocardial relaxation. Circ Res 111, 718–727, doi:10.1161/CIRCRESAHA.112.274464 (2012).

19 Robinson, P. et al. Hypertrophic cardiomyopathy mutations increase myofilament Ca(2+) buffering, alter intracellular Ca(2+) handling, and stimulate Ca(2+)-dependent signaling. J Biol Chem 293, 10487–10499, doi:10.1074/jbc.RA118.002081 (2018).

20 Toepfer, C. N. et al. Hypertrophic cardiomyopathy mutations in MYBPC3 dysregulate myosin. Sci Transl Med 11, doi:10.1126/scitranslmed.aat1199 (2019).

21 Hooijman, P., Stewart, M. A. & Cooke, R. A new state of cardiac myosin with very slow ATP turnover: a potential cardioprotective mechanism in the heart. Biophys J 100, 1969–1976, doi:10.1016/j.bpj.2011.02.061 (2011).

22 Burdyga, A. et al. Phosphatases control PKA-dependent functional microdomains at the outer mitochondrial membrane. Proc Natl Acad Sci U S A 115, E6497–E6506, doi:10.1073/pnas.1806318115 (2018).

23 Sikkel, M. B. et al. Hierarchical statistical techniques are necessary to draw reliable conclusions from analysis of isolated cardiomyocyte studies. Cardiovasc Res 113, 1743–1752, doi:10.1093/cvr/cvx151 (2017).

24 Van Sligtenhorst, I. et al. Cardiomyopathy in alpha-kinase 3 (ALPK3)-deficient mice. Vet Pathol 49, 131–141, doi:10.1177/0300985811402841 (2012).

25 Chandra, M. et al. Ca(2+) activation of myofilaments from transgenic mouse hearts expressing R92Q mutant cardiac troponin T. Am J Physiol Heart Circ Physiol 280, H705–713, doi:10.1152/ajpheart.2001.280.2.H705 (2001).

26 Najafi, A. et al. Selective phosphorylation of PKA targets after beta-adrenergic receptor stimulation impairs myofilament function in Mybpc3-targeted HCM mouse model. Cardiovasc Res 110, 200–214, doi:10.1093/cvr/cvw026 (2016).

27 Wijnker, P. J., Murphy, A. M., Stienen, G. J. & van der Velden, J. Troponin I phosphorylation in human myocardium in health and disease. Neth Heart J 22, 463–469, doi:10.1007/s12471-014-0590-4 (2014).

28 Rao, V. S. et al. N-terminal phosphorylation of cardiac troponin-I reduces length-dependent calcium sensitivity of contraction in cardiac muscle. J Physiol 591, 475–490, doi:10.1113/jphysiol.2012.241604 (2013).

29 Toepfer, C. N. et al. Myosin Sequestration Regulates Sarcomere Function, Cardiomyocyte Energetics, and Metabolism, Informing the Pathogenesis of Hypertrophic Cardiomyopathy. Circulation 141, 828–842, doi:10.1161/CIRCULATIONAHA.119.042339 (2020).

30 Sparrow, A. J., Watkins, H., Daniels, M. J., Redwood, C. & Robinson, P. Mavacamten rescues increased myofilament calcium sensitivity and dysregulation of Ca(2+) flux caused by thin filament hypertrophic cardiomyopathy mutations. Am J Physiol Heart Circ Physiol 318, H715–H722, doi:10.1152/ajpheart.00023.2020 (2020).

31 Green, E. M. et al. A small-molecule inhibitor of sarcomere contractility suppresses hypertrophic cardiomyopathy in mice. Science 351, 617–621, doi:10.1126/science.aad3456 (2016).

32 Zhao, M. et al. Gut microbiota production of trimethyl-5-aminovaleric acid reduces fatty acid oxidation and accelerates cardiac hypertrophy. Nat Commun 13, 1757, doi:10.1038/s41467-022-29060-7 (2022).

33 Blankenburg, R. et al. beta-Myosin heavy chain variant Val606Met causes very mild hypertrophic cardiomyopathy in mice, but exacerbates HCM phenotypes in mice carrying other HCM mutations. Circ Res 115, 227–237, doi:10.1161/CIRCRESAHA.115.303178 (2014).

34 Ehsan, M. et al. Mutant Muscle LIM Protein C58G causes cardiomyopathy through protein depletion. J Mol Cell Cardiol 121, 287–296, doi:10.1016/j.yjmcc.2018.07.248 (2018).

35 Vignier, N. et al. Nonsense-mediated mRNA decay and ubiquitin-proteasome system regulate cardiac myosin-binding protein C mutant levels in cardiomyopathic mice. Circ Res 105, 239–248, doi:10.1161/CIRCRESAHA.109.201251 (2009).

36 Schlossarek, S. et al. Adrenergic stress reveals septal hypertrophy and proteasome impairment in heterozygous Mybpc3-targeted knock-in mice. J Muscle Res Cell Motil 33, 5–15, doi:10.1007/s10974-011-9273-6 (2012).

37 Robinson, P. et al. Dilated cardiomyopathy mutations in thin-filament regulatory proteins reduce contractility, suppress systolic Ca(2+), and activate NFAT and Akt signaling. Am J Physiol Heart Circ Physiol 319, H306–H319, doi:10.1152/ajpheart.00272.2020 (2020).

38 Liu, Y., Chen, J., Fontes, S. K., Bautista, E. N. & Cheng, Z. Physiological and pathological roles of protein kinase A in the heart. Cardiovasc Res 118, 386–398, doi:10.1093/cvr/cvab008 (2022).

39 Schmid, M. & Toepfer, C. N. Cardiac myosin super relaxation (SRX): a perspective on fundamental biology, human disease and therapeutics. Biol Open 10, doi:10.1242/bio.057646 (2021).

40 Margara, F. et al. Mechanism based therapies enable personalised treatment of hypertrophic cardiomyopathy. Sci Rep 12, 22501, doi:10.1038/s41598-022-26889-2 (2022).

41 McNamara, J. W. et al. Ablation of cardiac myosin binding protein-C disrupts the super-relaxed state of myosin in murine cardiomyocytes. J Mol Cell Cardiol 94, 65–71, doi:10.1016/j.yjmcc.2016.03.009 (2016).

42 Toepfer, C. N. et al. SarcTrack. Circ Res 124, 1172–1183, doi:10.1161/CIRCRESAHA.118.314505 (2019).

43 Anderson, R. L. et al. Deciphering the super relaxed state of human beta-cardiac myosin and the mode of action of mavacamten from myosin molecules to muscle fibers. Proc Natl Acad Sci U S A 115, E8143–E8152, doi:10.1073/pnas.1809540115 (2018).

## References cited in SI

[s1] K. Gehmlich, M.S. Dodd, J.W. Allwood, M. Kelly, M. Bellahcene, H.V. Lad, A. Stockenhuber, C. Hooper, H. Ashrafian, C.S. Redwood, L. Carrier, W.B. Dunn, Changes in the cardiac metabolome caused by perhexiline treatment in a mouse model of hypertrophic cardiomyopathy, Mol Biosyst 11(2) (2015) 564–73.

[s2] A.N. Abou Tayoun, T. Pesaran, M.T. DiStefano, A. Oza, H.L. Rehm, L.G. Biesecker, S.M. Harrison, G. ClinGen Sequence Variant Interpretation Working, Recommendations for interpreting the loss of function PVS1 ACMG/AMP variant criterion, Hum Mutat 39(11) (2018) 1517-1524.

[s3] S. Richards, N. Aziz, S. Bale, D. Bick, S. Das, J. Gastier-Foster, W.W. Grody, M. Hegde, E. Lyon, E. Spector, K. Voelkerding, H.L. Rehm, A.L.Q.A. Committee, Standards and guidelines for the interpretation of sequence variants: a joint consensus recommendation of the American College of Medical Genetics and Genomics and the Association for Molecular Pathology, Genet Med 17(5) (2015) 405–24.

