## Supplementary material for "An *ALPK3* truncation variant causing autosomal dominant hypertrophic cardiomyopathy is partially rescued by mavacamten": Electronic Supporting Information

#### Supplemental material and methods

##### *Alpk3 knock-out mice*

*Alpk3* knock-out (KO) mice (Alpk3$^{tm1b(EUCOMM)Hmgu}$) were obtained from EUCOMM, embryo-rederived and backcrossed onto C57BL/6JOlaHsd (Envigo RMS (UK) Ltd) for at least 6 generations until congenic before phenotyping.

Genotyping was performed using the REDExtract-N-Amp™ Tissue PCR Kit (Sigma, for primer pairs see below) and separate PCRs for the presence of the WT and the KO allele. PCR products were analysed by agarose electrophoresis (1.5 % agarose in Tris-Borate-EDTA buffer, Fisher).

*WT allele (401 bp product)*

Forward primer: 5’ – GGTCAAGACTCCATTCAGCCCT – 3’

Reverse primer: 5’ – CCTCCACGCCTATTCTAGCCTC – 3’

*KO allele (358 bp product):*

Forward primer: 5’ – GCGAGCTCAGACCATAACTT – 3’

Reverse primer: 5’ – CCCAAGTCACAAACTGTTCA – 3’

##### *Quantitative Reverse Transcriptase Polymerase Chain Reaction (qPCR)*

qPCRs were performed as described [1] using the Taqman probes (Applied Biosystems) listed in Table S7.

***Cell measurements***

Cardiomyocytes were isolated from the three genotypes and fixed (see Immunofluorescence). Individual cells were documented on bright field setting with an EVOS M5000 imaging system (Thermo), using a 20x objective. In ImageJ, length and width of individual rod-shaped cardiomyocytes were determined manually for each cell and recorded. For statistical analysis, ROUT outlier test was performed and outliers (1/118 in Het and 1/163 in Hom for length and 12/118 for Het and 1/163 for Hom for width omitted). Data was analysed with nested 1-way ANOVA taking into consideration that cells were derived from 2 WT, 3 Het and 3 Hom mice.

**Supplemental tables and figures**

**Table S1:** *ALPK3tv* detected in NIHR Bioresource Rare Disease HCM project (BRRD) cohort (HCM) and in GnomAD (controls).

**Table S2:** HCM case versus control analyses

**Abbreviations** echocardiography parameters

**Table S3.** Summary of echocardiography parameters of wildtype (WT), heterozygous (Het) and homozygous (Hom) *Alpk3 K201X* mice aged 3 months as well as WT and Het at 6 months.

**Table S4.** Summary of echocardiography parameters of homozygous global *Alpk3* knock out mice (KO) versus homozygous *Alpk3* K201X mice at 3 months.

**Table S5.** Summary of echocardiography parameters of wildtype (WT) and heterozygous (Het) *Alpk3 K201X* mice after chronic adrenergic challenge (Iso/PE) or sham saline treatment.

**Table S6.** Extracted parameters for unloaded sarcomere shortening and $Ca^{2+}$ transient measurements from isolated left ventricular cardiomyocytes.

**Table S7.** Taqman Assays for qPCR

**Figure S1.** Position of *ALPK3tv* in our HCM cohort and generation of a novel mouse model *Alpk3* K201X.

**Figure S2.** Morphological and functional comparison of the hearts

**Figure S3.** qPCR demonstrates induction of transcripts related to foetal gene programme and hypertrophic signalling in homozygous (Hom) *Alpk3* K201X Hom mice.

**Figure S4.** Cellular hypertrophy, but no evidence of fibrosis in hearts of *Alpk3* K201X mice

**Figure S5.** *Alpk3* K201X mice show no evidence for dysregulated myomesin

**Figure S6.** Isolated left ventricular cardiomyocytes from *Alpk3* K201X mice in the absence of the $Ca^{2+}$ indicator fura2 display hypercontractile unloaded sarcomere shortening with prolonged relaxation.

**Figure S7.** Cluster analysis illustration from all significant extracted parameters taken from unloaded sarcomere shortening and calcium transient measurements.

**Figure S8.** Whole western blot images.

**Table S1.** ***ALPK3tv*** **detected in NIHR Bioresource Rare Disease HCM project (BRRD) cohort (HCM) and in GnomAD (controls).** Genomic co-ordinates according to genome buildGRCh38. Coding (c.) and protein (p.) nomenclature according to MANE transcripts (NM_929778.5; ENST0000025888.6). ACMG criteria and classification according to [2, 3]. * Variants in last exon in the gene, may escape nonsense-mediated decay.

| **Genomic co-ordinates (chromosome 15)** | **c.** | **p.** | **Predicted effect** | **HCM/control** | **ACMG criteria** | **Classification** |
|---|---|---|---|---|---|---|
| g.84817514del | c.62del | p.(Gly21Alafs*88) | Frameshift | HCM | PVS1 | Likely pathogenic |
| g.84839880A>T | c.601A>T | p.(Lys201*) | Nonsense | HCM | PVS1; PM2 | Likely pathogenic |
| g.84840483del | c.1204del | p.(Ala402Profs*14) | Frameshift | HCM | PVS1; PM2 | Likely pathogenic |
| g.84856391G>A | c.1654-1G>A | p.? | Splice site | HCM | PVS1; PM2 | Likely pathogenic |
| g.84862795C>G | c.4290C>G | p.(Tyr1430*) | Nonsense | HCM | PVS1; PM2 | Likely pathogenic |
| g.84839076del | c.401del | p.(Arg134Profs*39) | Frameshift | Control | PVS1; PM2 | Likely pathogenic |
| g.84840372C>T | c.1093C>T | p.(Gln365*) | Nonsense | 2 x Controls | PVS1 | Likely pathogenic |
| g.84856782G>T | c.2044G>T | p.(Glu682*) | Nonsense | Control | PVS1 | Likely pathogenic |
| g.84857193C>T | c.2455C>T | p.(Arg819*) | Nonsense | Control | PVS1 | Likely pathogenic |
| g.84862718C>T | c.4213C>T | p.(Arg1405*) | Nonsense | Control | PVS1 | Likely pathogenic |
| g.84868446del | c.5108del | p.(Gly1703Alafs*53) | Frameshift** | Control | PVS1_moderate; PM2 | VUS |
| g.84868453del | c.5115del | p.(*1706Serext*49) | Stop loss** | 2 x Controls | PVS1_moderate | VUS |

**Table S2. HCM case versus control analyses** using 230 HCM cases and 6,219 controls (see Material and Methods). FET=Fishers exact test. OR=Odds ratio. Lower CI=lower 95% confidence interval of OR. Upper CI=upper 95% Confidence interval of OR.

| Variant type | HCM case frequency | Control frequency | FET p-value | OR | Lower_CI | Upper CI |
|---|---|---|---|---|---|---|
| **All variants** | 0.0565 | 0.0263 | 0.0116 | 2.2147 | 1.1394 | 3.9596 |
| **Missense** | 0.0348 | 0.0243 | 0.2788 | 1.45 | 0.61 | 2.97 |
| **Truncating** | 0.0217 | 0.0014 | **0.0001** | **16.07** | 4.27 | 52.05 |
| **Non-truncating** | 0.0348 | 0.0250 | 0.3869 | 1.41 | 0.59 | 2.88 |

**Abbreviations echocardiography parameters**

| | |
|---|---|
| **EF (%)** | Ejection fraction |
| **FS SAX (%)** | Fractional Shortening in short axis view |
| **LV Mass (mg)** | Left ventricle mass calculated by system |
| **LVAW:d (mm)** | Left anterior wall thickness in diastole |
| **LVAW:s (mm)** | Left anterior wall thickness in systole |
| **LVPW:d (mm)** | Left ventricular posterior wall thickness in diastole |
| **LVPW:s (mm)** | Left ventricular posterior wall thickness in systole |
| **LVID:d (mm)** | Left ventricular internal diameter in diastole |
| **LVID:s (mm)** | Left ventricular internal diameter in systole |
| **LV Vol:d (uL)** | Left ventricular volume in diastole |
| **LV Vol:s (uL)** | Left ventricular volume in systole |
| **PSV (mm/sec)** | Peak systolic velocity |
| **HR (/min)** | Heart rate |
| **HW/TL (g/mm)** | Heart weight to tibia length |

**Table S3. Summary of echocardiography parameters of wildtype (WT), heterozygous (Het) and homozygous (Hom) *Alpk3 K201X* mice aged 3 months as well as WT and Het at 6 months.**

Data shown as mean ± SEM. Where not all animals of a genotype could be analysed, the number of animals analysed for this parameter stated in brackets. Statistical significance indicated as for Hom versus WT at 3 months; ** p < 0.01, *** < p<0.001, **** p < 0.001; Kruskal-Wallis test. No significant changes were observed for Het versus WT at either age (Student's t-test at 6 months).

| **Age** | **3 months** | | | | **6 months** | |
|---|---|---|---|---|---|---|
| | **WT** | **Het** | **Hom** | | **WT** | **Het** |
| **N** | 23 | 21 | 8 | | 10 | 9 |
| **EF (%)** | 66±2 | 61±2 | 28±3 **** | | 67±3 | 66±4 |
| **FS SAX (%)** | 36.5±1.7 | 37±3 | 12.9±1.5 **** | | 37± 2 | 37±3 |
| **LV Mass (mg)** | 117±7 | 124±6 | 157±8 ** | | 118±5 | 125±9 |
| **LVAW:d (mm)** | 0.91±0.05 | 0.95±0.04 | 1.01±0.07 | | 0.99±0.05 | 1.07±0.09 |
| **LVAW:s (mm)** | 1.38±0.07 | 1.34±0.07 | 1.18±0.09 | | 1.24±0.06 | 1.40±0.08 |
| **LVPW:d (mm)** | 0.99±0.06 | 0.91±0.05 | 1.06±0.05 | | 0.90±0.05 | 0.88±0.03 |
| **LVPW:s (mm)** | 1.45±0.05 (20) | 1.29±0.07 (18) | 1.25±0.04 | | 1.4±0.06 | 1.36±0.11 |
| **LVID:d (mm)** | 3.91±0.07 | 4.03±0.09 | 4.42±0.09 ** | | 3.970±0.07 | 4.00±0.10 |
| **LVID:s (mm)** | 2.50±0.10 | 2.73±0.11 | 3.86±0.11**** | | 2.50± 0.12 | 2.55±0.17 |
| **LV Vol:d (uL)** | 67±3 | 72±4 | 89±4 ** | | 69±3 | 71±4 |
| **LV Vol:s (uL)** | 24±2 | 29±3 | 65±4 **** | | 23 ±3 | 25±3 |
| **PSV (mm/sec)** | 19.8±0.9 (22) | 19.6±0.9 | 12.2±1.1 *** | | 21.4±0.9 | 20.7±0.5 |
| **HR (/min)** | 478±6 (20) | 469±5 | 488±4 | | 466±13 | 486±12 |
| **HW/TL (g/mm)** | 7.4±0.3 (16) | 8.5±0.3 (9) | 13.1±0.2 **** (16) | | 8.2±0.2 | 8.9±0.3 |

**Table S4. Summary of echocardiography parameters of homozygous global *Alpk3* knock out mice (KO) versus homozygous *Alpk3* K201X mice at 3 months.**

Data shown as mean ± SEM. Number of animals per genotype and parameter stated in brackets. No statistical differences in the phenotypes of the two models were observed, apart from minor difference in heart rate (* $p < 0.05$; Student's t-test used).

Please note the *Alpk3* K201X (Hom) cohort is a different cohort to the one shown in Fig. 1 and Table S3.

| | ***Alpk3* KO** | ***Alpk3* K201X (Hom)** |
|---|---|---|
| **N** | 13 | 13 |
| **EF (%)** | 30.6±1.8 | 36±4 |
| **FS SAX (%)** | 14.4±0.9 | 15.6±1.4 (12) |
| **LV Mass (mg)** | 166±7 | 177±9 |
| **LVAW:d (mm)** | 1.03±0.04 | 1.12±0.05 |
| **LVAW:s (mm)** | 1.14±0.03 | 1.16±0.05 |
| **LVPW:d (mm)** | 1.03±0.04 | 1.05±0.05 |
| **LVPW:s (mm)** | 1.17±0.04 | 1.33±0.09 |
| **LVID:d (mm)** | 4.57±0.09 | 4.55±0.08 |
| **LVID:s (mm)** | 3.92±0.10 | 3.76±0.13 |
| **LV Vol:d (uL)** | 97±5 | 95±4 |
| **LV Vol:s (uL)** | 67±4 | 62±4 |
| **HR (/min)** | 471±3 | 480±3 * |
| **HW/TL (mg/mm)** | 12.6±0.4 | 13.3±0.3 |

**Table S5. Summary of echocardiography parameters of wildtype (WT) and heterozygous (Het) *Alpk3 K201X* mice after chronic adrenergic challenge (Iso/PE) or sham saline treatment.**

Data shown as mean ± SEM. Statistical significance indicated as * $p < 0.05$, 2-way-ANOVA test with Tukey test for multiple comparisons used. Sample size (N) in table refers to animals per group, for HW/TL n numbers are stated in brackets if different.

$^{\$}$ $p < 0.05$ versus WT Iso/PE,

* $p < 0.05$, ** $p < 0.01$, *** $p < 0.001$, **** $p < 0.0001$ versus Het sham,

$^{\#}$ $p < 0.05$, $^{\#\#\#}$ $p < 0.001$ versus WT sham

| | sham | | Iso/PE | |
|---|---|---|---|---|
| | **WT** | **Het** | **WT** | **Het** |
| **N** | 12 | 8 | 10 | 12 |
| **EF (%)** | 67±5 | 65±5 | 72±4 | 68±3 |
| **FS SAX (%)** | 38±3 | 36±3 | 41±3 | 38±2 |
| **LV Mass (mg)** | 104±6 | 108±8 | 125±6 | 157±8***,$^{\$}$ |
| **LVAW:d (mm)** | 0.96±0.05 | 0.86±0.05 | 1.02±0.05 | 1.17±0.05*** |
| **LVAW:s (mm)** | 1.24±0.06 | 1.24±0.07 | 1.53±0.08$^{\#}$ | 1.60±0.07** |
| **LVPW:d (mm)** | 0.95±0.04 | 0.99±0.08 | 1.03±0.04 | 1.06±0.04 |
| **LVPW:s (mm)** | 1.41±0.10 | 1.37±0.07 | 1.58±0.07 | 1.53±0.06 |
| **LVID:d (mm)** | 3.65±0.09 | 3.84±0.11 | 3.84±0.14 | 4.10±0.12 |
| **LVID:s (mm)** | 2.28±0.15 | 2.48±0.17 | 2.28±0.19 | 2.55±0.12 |
| **LV Vol:d (uL)** | 57± 3 | 64±4 | 65±5 | 75±5 |
| **LV Vol:s (uL)** | 19±3 | 23±4 | 20±4 | 24±3 |
| **PSV (mm/sec)** | 20.2±0.7 | 21.1±1.9 | 21.8±1.5 | 22.7±1.2 |
| **HR(/min)** | 482±4 | 472±6 | 539±26 | 549±24* |
| **HW/TL (mg/mm)** | 8.4±0.3 | 8.7±0.3 (10) | 9.97±0.16$^{\#\#\#}$ (11) | 11.1±0.3****,$^{\$}$ (10) |

**Table S6. Extracted parameters** for (**A**) unloaded sarcomere shortening (fura2 loaded), (**B**) $Ca^{2+}$ transient, and (**C**) indicator free unloaded sarcomere shortening measurements from left ventricular cardiomyocytes isolated from WT, heterozygous (Het) and homozygous (Hom) *Alpk3* K201X cardiomyocytes, with and without treatment with 0.5 μM mavacamten.

Integers are ±SEM, bold text indicates significance $p < 0.05$. * $p < 0.05$, ** $p < 0.01$, *** $p < 0.001$, **** $p < 0.0001$ using nested 1-way-ANOVA to adjust for hierarchical clustering of individual data sets between mice for (**A**) and (**B**) and Kruskal-Wallis with Dunn's correction for (**C**). Exact sample size (n) per condition (cells per group of mice) is given in the table.

## A

| | (*n*) mice | (*n*) cells | Basal Sarcomere Length (μm) | Peak Sarcomere Length (μm) | Peak Height (μm) | Fractional Shortening (%) | T Peak Contraction (sec) | T10% Contraction (sec) | T50% Contraction (sec) | T90% contraction (sec) | T10% Relaxation (sec) | T50% Relaxation (sec) | T90% Relaxation (sec) |
|---|---|---|---|---|---|---|---|---|---|---|---|---|---|
| WT | 6 | 157 | 1.877±0.006 | 1.787±0.007 | 0.090±0.004 | 4.779±0.195 | 0.046±0.001 | 0.010±0.001 | 0.021±0.001 | 0.034±0.001 | 0.011±0.001 | 0.028±0.001 | 0.072±0.003 |
| WT + 0.5 μM mava | 6 | 164 | **1.955±0.004 ****** | **1.920±0.005 ****** | **0.035±0.002 ****** | **1.802±0.109 ****** | 0.042±0.001 | 0.011±0.001 | 0.021±0.002 | 0.033±0.001 | 0.011±0.001 | 0.029±0.001 | 0.071±0.004 |
| Het | 8 | 192 | **1.795±0.005 ****** | **1.727±0.008 ****** | **0.068±0.003 **** | **3.788±0.166 **** | 0.055±0.003 | 0.014±0.001 | 0.025±0.001 | 0.041±0.001 | 0.014±0.001 | 0.035±0.001 | **0.096±0.003 *** |
| Het + 0.5 μM mava | 8 | 182 | 1.901±0.004 | **1.872±0.005 ****** | **0.029±0.002 ****** | **1.540±0.094 ****** | 0.050±0.002 | 0.013±0.001 | 0.024±0.001 | 0.038±0.002 | 0.012±0.001 | 0.031±0.001 | 0.085±0.003 |
| Hom | 5 | 164 | **1.736±0.005 ****** | **1.663±0.005 ****** | 0.074±0.003 | 4.065±0.158 | **0.084±0.002 ****** | 0.017±0.001 | **0.033±0.001 ***** | **0.059±0.001 ****** | **0.027±0.001 *** | **0.065±0.002 ****** | **0.134±0.003 ****** |
| Hom + 0.5 μM mava | 5 | 167 | 1.850±0.005 | **1.829±0.005 **** | **0.022±0.001 ****** | **1.165±0.074 ****** | **0.069±0.003 *** | 0.018±0.001 | **0.033±0.002 ***** | **0.053±0.003 *** | 0.019±0.001 | **0.055±0.002 ***** | **0.126±0.004 ****** |

## B

| | (*n*) mice | (*n*) cells | Diastolic $_i[Ca^{2+}]$ (μM) | Systolic $_i[Ca^{2+}]$ (μM) | $Ca^{2+}$ Transient Amplitude (μM) | T Peak Release (sec) | T10% $Ca_{2+}$ Release (sec) | T50% $Ca_{2+}$ Release (sec) | T90% $Ca_{2+}$ Release (sec) | T10% $Ca_{2+}$ Reuptake (sec) | T50% $Ca_{2+}$ Reuptake (sec) | T90% $Ca_{2+}$ Reuptake (sec) |
|---|---|---|---|---|---|---|---|---|---|---|---|---|
| WT | 6 | 157 | 0.483±0.020 | 1.485±0.041 | 1.002±0.032 | 0.025±0.001 | 0.005±0.001 | 0.009±0.001 | 0.015±0.001 | 0.021±0.001 | 0.069±0.001 | 0.170±0.003 |
| WT + 0.5 μM mava | 6 | 164 | 0.332±0.018 | 1.237±0.035 | 0.905±0.030 | 0.025±0.001 | 0.006±0.001 | 0.008±0.001 | 0.015±0.002 | 0.024±0.001 | 0.068±0.001 | 0.156±0.003 |
| Het | 8 | 192 | **0.719±0.026 ***** | **1.963±0.064 *** | 1.244±0.047 | 0.030±0.001 | 0.009±0.001 | 0.011±0.001 | 0.019±0.001 | 0.023±0.001 | 0.076±0.001 | 0.186±0.003 |
| Het + 0.5 μM mava | 8 | 182 | 0.464±0.021 | 1.506±0.045 | 1.042±0.035 | 0.028±0.001 | 0.007±0.001 | 0.010±0.001 | 0.017±0.001 | 0.022±0.001 | 0.066±0.001 | 0.161±0.003 |
| Hom | 5 | 164 | 0.798±0.027 | **2.067±0.051 **** | 1.269±0.038 | **0.034±0.001 *** | 0.008±0.001 | 0.011±0.001 | **0.021±0.001 *** | **0.032±0.001 ***** | **0.091±0.001 **** | 0.192±0.003 |
| Hom + 0.5 μM mava | 5 | 167 | **0.442±0.024 ***** | 1.261±0.036 | 0.819±0.032 | 0.033±0.001 | 0.008±0.001 | 0.013±0.001 | 0.022±0.001 | **0.033±0.001 ***** | **0.089±0.001 *** | 0.181±0.003 |

## C

| | (*n*) mice | (*n*) cells | Basal Sarcomere Length (μm) | Peak Sarcomere Length (μm) | Peak Height (μm) | Fractional Shortening (%) | T Peak Contraction (sec) | T10% Contraction (sec) | T50% Contraction (sec) | T90% contraction (sec) | T10% Relaxation (sec) | T50% Relaxation (sec) | T90% Relaxation (sec) |
|---|---|---|---|---|---|---|---|---|---|---|---|---|---|
| WT | 2 | 35 | 1.872±0.014 | 1.758±0.015 | 0.114±0.006 | 6.779±0.312 | 0.051±0.002 | 0.012±0.001 | 0.024±0.001 | 0.039±0.001 | 0.011±0.001 | 0.026±0.001 | 0.059±0.003 |
| WT + 0.5 μM mava | 2 | 35 | 1.922±0.013 | **1.858±0.013 ****** | **0.066±0.008 ****** | **3.283±0.319 ****** | 0.050±0.003 | 0.014±0.001 | 0.022±0.002 | 0.035±0.002 | 0.013±0.001 | 0.037±0.002 | 0.068±0.005 |
| Het | 1 | 34 | **1.806±0.010 ****** | **1.716±0.008 ****** | 0.091±0.003 | 6.910±0.321 | **0.064±0.004 *** | 0.016±0.001 | 0.031±0.001 | 0.044±0.001 | **0.025±0.001 *** | **0.057±0.004 **** | **0.095±0.004 ****** |
| Het + 0.5 μM mava | 1 | 15 | 1.926±0.010 | **1.872±0.015 ****** | **0.054±0.007 ****** | **2.790±0.368 ****** | 0.052±0.003 | 0.013±0.001 | 0.025±0.002 | 0.038±0.002 | 0.012±0.002 | 0.030±0.003 | 0.071±0.005 |
| Hom | 2 | 46 | **1.721±0.015 ****** | **1.656±0.014 ****** | **0.065±0.006 ****** | **3.765±0.321 ****** | **0.066±0.002 ***** | 0.014±0.001 | 0.029±0.001 | **0.049±0.002 ***** | **0.018±0.001 *** | **0.047±0.003 ***** | **0.109±0.005 ****** |
| Hom + 0.5 μM mava | 2 | 39 | 1.878±0.011 | **1.836±0.012 ****** | **0.042±0.004 ****** | **2.253±0.191 ****** | **0.071±0.003 ****** | 0.016±0.001 | **0.032±0.002 *** | **0.052±0.003 ***** | **0.021±0.001 *** | **0.054±0.003 ***** | **0.124±0.006 ****** |

**Table S7. Taqman Assays for qPCR** (all Applied Biosystems, FAM-MGB unless stated otherwise)

| *Transcript* | *Species* | *Assay ID* | *Comments* |
|---|---|---|---|
| Acta1 | Mouse | Mm00808218_g1 | |
| Alpk3 | Mouse | Mm00475369_m1 | |
| Ankrd1 | Mouse | Mm00496512_m1 | |
| Ankrd2 | Mouse | Mm00508030_m1 | |
| Col1a1 | Mouse | Mm00801666_g1 | |
| Fhl1 | Mouse | Mm04204611_g1 | |
| Gapdh | Mouse | 4352339E | VIC-MGB labelled |
| Myh7 | Mouse | Mm00600555_m1 | |
| Nppa | Mouse | Mm01255748_g1 | |
| Nppb | Mouse | Mm01255770_g1 | |
| Rcan1.4 | Mouse | Mm00627762_m1 | |

**Figure S1. Position of *ALPK3tv* in our HCM cohort and generation of a novel mouse model *Alpk3* K201X.**

(**A**) **Positions of the heterozygous *ALPK3tv***, identified in adult onset HCM patients. Coding exons are numbered and positions of domains are indicated by colour. *ALPK3* K201X is printed in bold.

(**B**) **Sanger sequencing traces of the mouse model.** Top trace from a wildtype (WT) mouse and bottom trace from heterozygous *Alpk3* K201X mouse (Het) are shown. The codon 201 is indicated in blue (AAG coding for lysine, Lys, in WT and TAG coding for a Stop codon in Het). Two silent modifications are also indicated: one creates a *HindIII* restriction site in the Het, the other disrupts the PAM sequence.

(**C**) **Transcript levels of *Alpk3* in the novel mouse model *Alpk3* K201X.** *Alpk3* transcript was measured in WT (blue), *Alpk3* K201X Het (red) and *Alpk3* K201X Hom (grey) and Alpk3 knockout (KO, white) hearts relative to *Gapdh* by qPCR. Data was not normally distributed (Shapiro-Wilk test), hence Kruskal-Wallis test with Dunn's post hoc test was used. *Alpk3* was found to be reduced in Hom and KO mice with * $p < 0.05$ and *** $p < 0.001$, respectively. N numbers: WT 6, Het 6, Hom 5, KO 6.

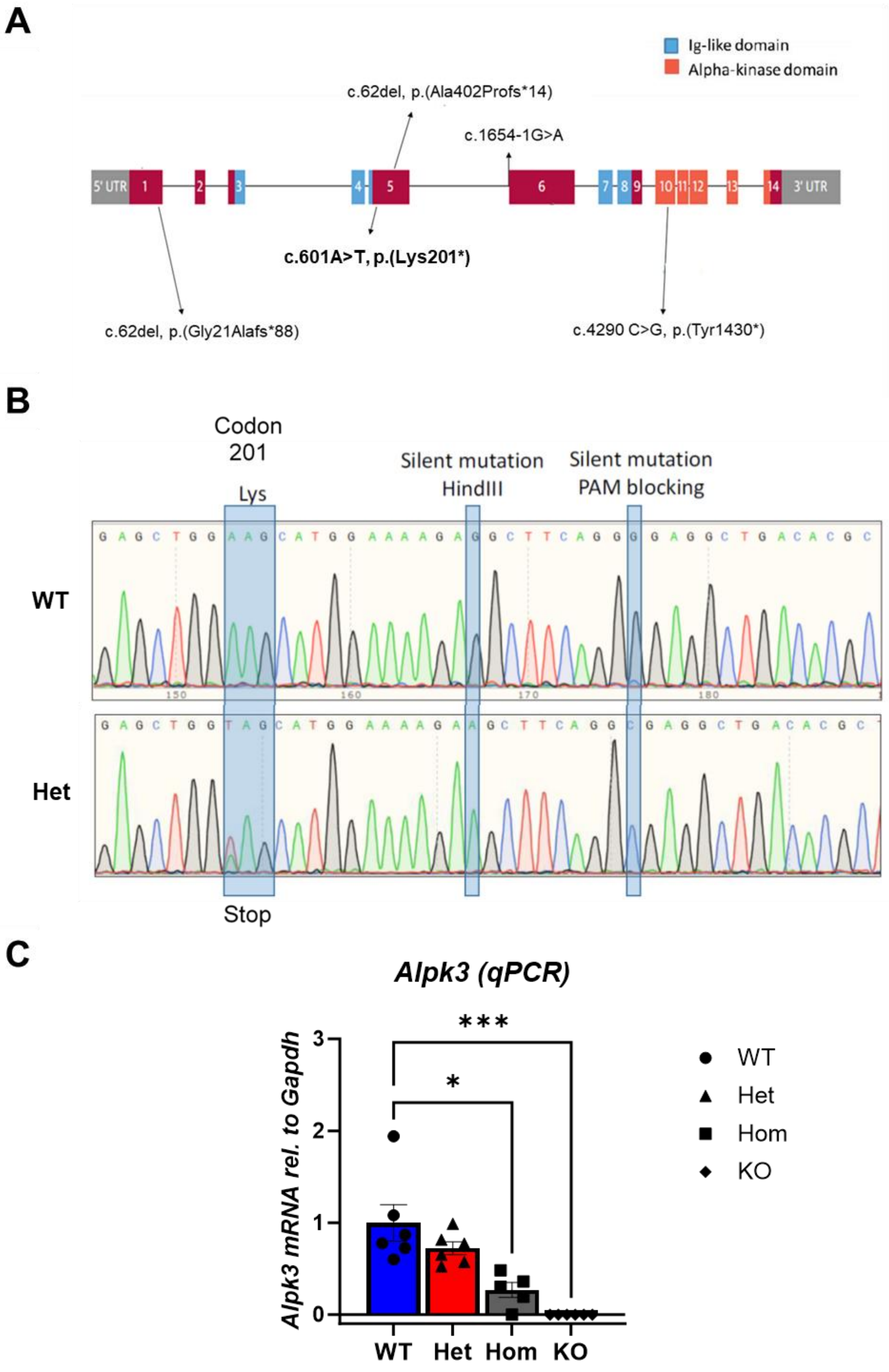
A
Ig-like domain
Alpha-kinase domain
c.62del, p.(Ala402Profs*14)
c.1654-1G>A
5' UTR
1
2
3
4
5
6
7
8
9
10
11
12
13
14
3' UTR
c.601A>T, p.(Lys201*)
c.62del, p.(Gly21Alafs*88)
c.4290 C>G, p.(Tyr1430*)
B
Codon 201
Lys
Silent mutation HindIII
Silent mutation PAM blocking
WT
Het
Stop
C
Alpk3 (qPCR)
Alpk3 mRNA rel. to Gapdh
***
*
WT
Het
Hom
KO

**Figure S2. Morphological and functional comparison of the hearts**

(**A**) Representative images of haematoxylin-eosin stained, long-axis heart sections of WT, *Alpk3* K201X Het and *Alpk3* K201X Hom mice to demonstrate the cardiac morphology.

(**B**) Representative examples of echocardiography (M-Mode) of WT, *Alpk3* K201X Het and *Alpk3* K201X Hom.

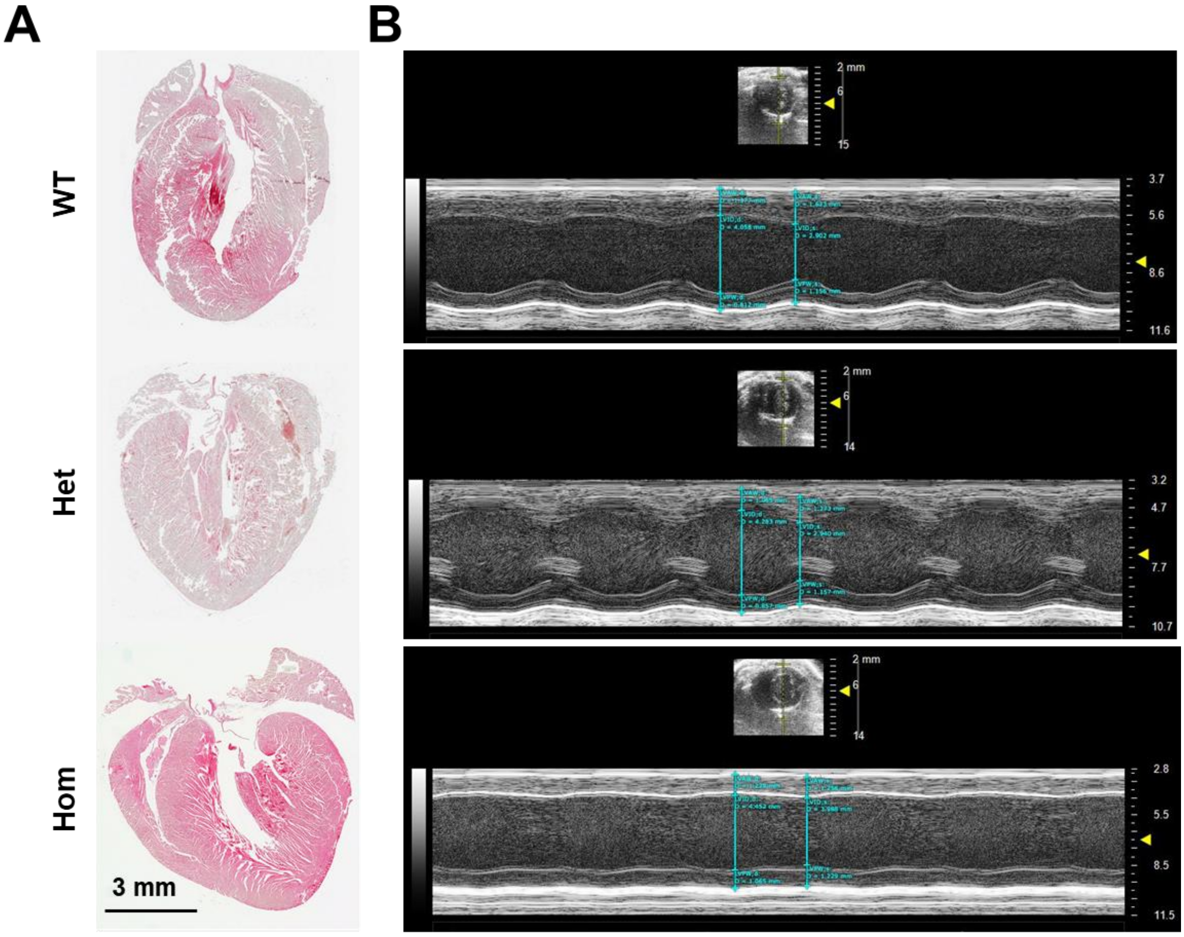

**Figure S3. qPCR demonstrates induction of transcripts related to foetal gene programme and hypertrophic signalling in homozygous (Hom) *Alpk3* K201X mice.**

Transcripts related to the fetal gene programme (**A**) and hypertrophic signalling (**B**) were measured in WT (blue), *Alpk3* K201X Het (red) and *Alpk3* K201X Hom (grey) and Alpk3 knockout (KO, white) hearts relative to Gapdh by qPCR. No induction as observed for Het mice, while induction of foetal gene programme and hypertrophic signalling was observed for Hom and KO mice.

Data was normally distributed (Shapiro-Wilk test) for all but *Ankrd1* and *Fhl1*. For normally distributed data, 1-way-ANOVA with Dunnett's post hoc test was applied; for Ankrd1 and Fhl1 Kruskal Wallis test with Dunn's post hoc test was used. * $p < 0.05$, ** $p < 0.01$, *** $p < 0.001$, **** $p < 0.0001$. N numbers: (A) WT 5, Het 6, Hom 4, KO 6; (B) WT 6, Het 6, Hom 5, KO 6.

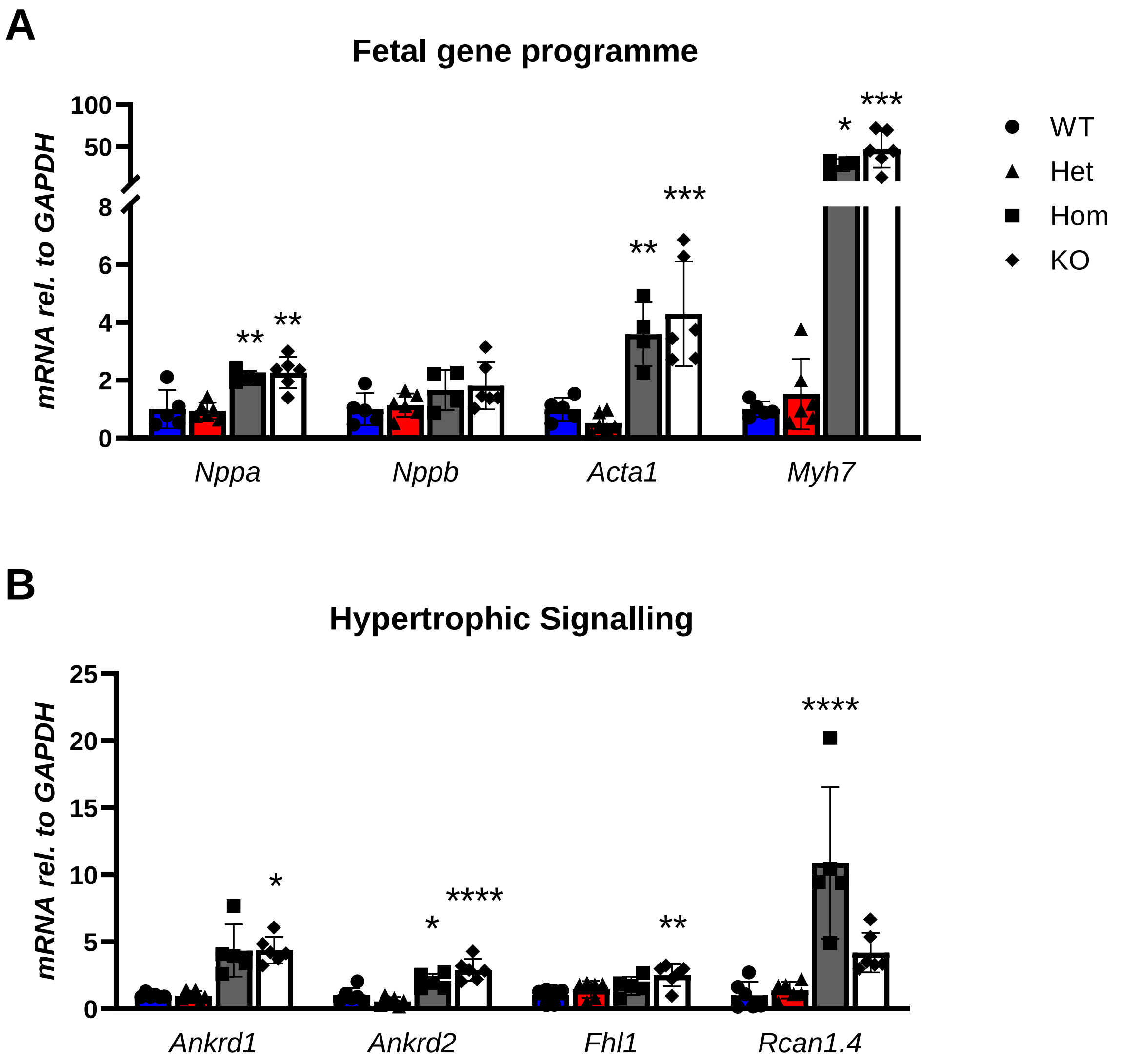

**Figure S4. Cellular hypertrophy, but no evidence of fibrosis in hearts of *Alpk3* K201X mice**

(**A**) **Histology of *Alpk3* K201X is unremarkable.** Representative images of haematoxylin-eosin stained, cardiac sections of wild type (WT), heterozygous (Het) *Alpk3* K201X and homozygous (Hom) *Alpk3* K201X mice shows no striking abnormalities. The same is true for homozygous *Alpk3* knockout mice (KO). Scale bare represents 50 microns.

(**B**) **No molecular evidence of fibrosis in heart *Alpk3* K201X mice.** Collagen *Col1a1* transcript was measured in WT (blue), *Alpk3* K201X Het (red) and *Alpk3* K201X Hom (grey) and Alpk3 knockout (KO, white) hearts relative to *Gapdh* by qPCR. Data was not normally distributed (Shapiro-Wilk test), hence Kruskal-Wallis test with Dunn's post hoc test was used. No evidence of induction of *Col1a1* was found in any of the genotypes, suggesting no induction of fibrosis. N numbers: WT 6, Het 6, Hom 5, KO 6.

(**C**) **Increased length, but not width in cardiomyocytes isolated from *Alpk3* K201X mice.** Isolated fixed cardiomyoctes from WT (blue), heterozygous (Het, red) and homozygous (Hom, grey) *Alpk3* K201 mice were measured for length and width. Length was increased in Het and Hom cells, * $p < 0.05$, *** $p < 0.001$, nested 1-way ANOVA. 86 WT cells from 2 mice, 117 Het cells from 3 mice and 162 cells from 3 mice were measured for length; 86 WT cells from 2 mice, 106 Het cells from 3 mice and 162 cells from 3 mice were measured for width.

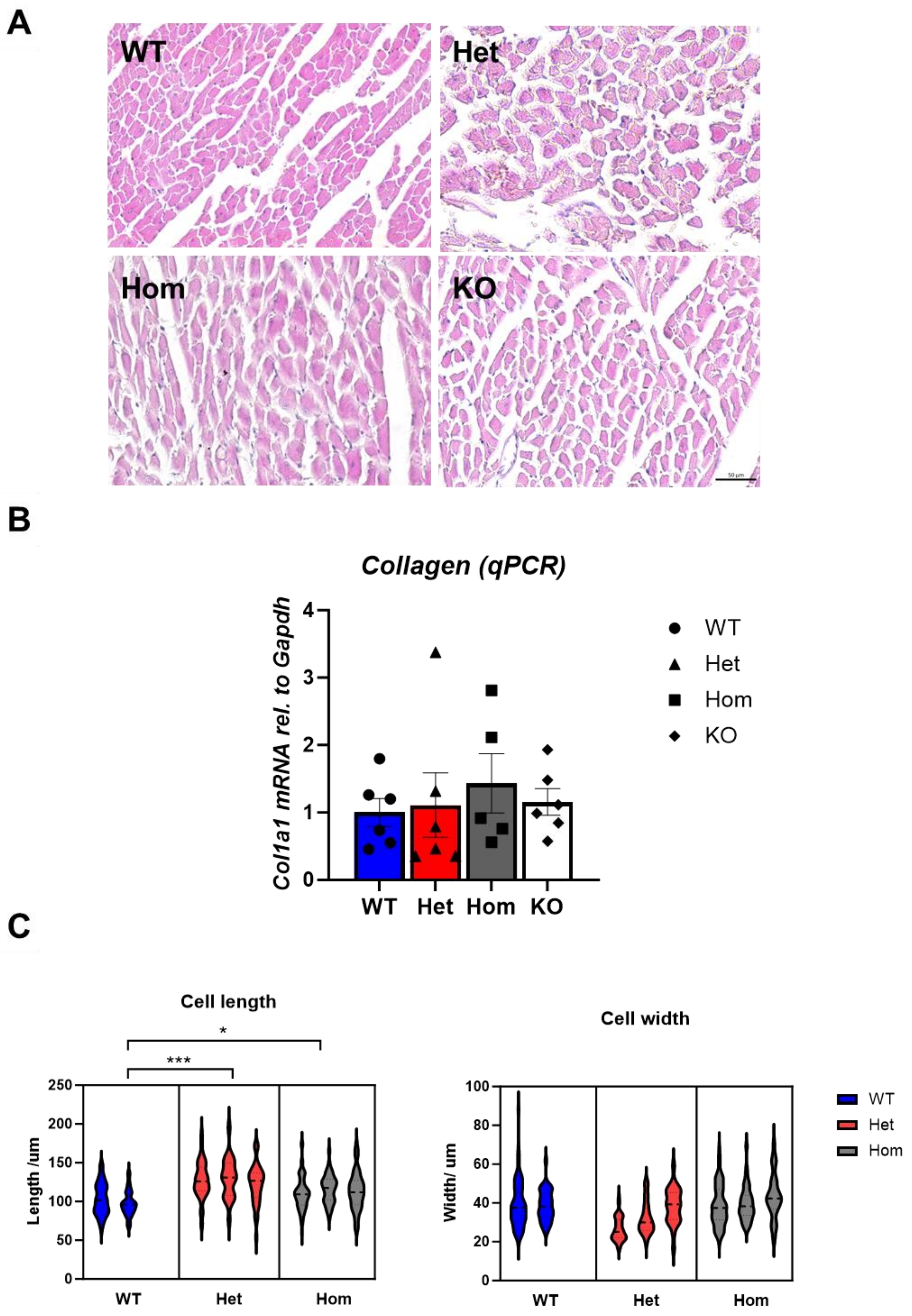
A
WT
Het
Hom
KO
B
Collagen (qPCR)
Col1a1 mRNA rel. to Gapdh
0
1
2
3
4
WT
Het
Hom
KO
C
Cell length
Length /um
0
50
100
150
200
250
*
***
WT
Het
Hom
Cell width
Width/ um
0
20
40
60
80
100
WT
Het
Hom

**Figure S5.** ***Alpk3*** **K201X mice show no evidence for dysregulated myomesin**

(**A**) Cardiac tissue of heterozygous (Het) and homozygous (Hom) for *Alpk3 K201X* as well as a wildtype (WT) control was stained for M-band protein myomesin (green), alpha-actinin 2 (red) and nuclei (NucBlue, blue). Representative immunofluorescence images of each genotype are shown. Scale bar equals 25 µm and 10 µm for inset. Sections from at least three animals per genotype (n = 3) were processed.

(**B**) Isolated cardiomyocytes of Het and Hom *Alpk3 K201X* mice well as a WT cells were stained for myomesin and alpha-actinin as above (but without visualisation of nuclei). Scale bar equals 10 µm. Slides with isolated cells from two animals per genotype (n=2) were processed.

No changes of myomesin such as accumulation or abundance were detectable.

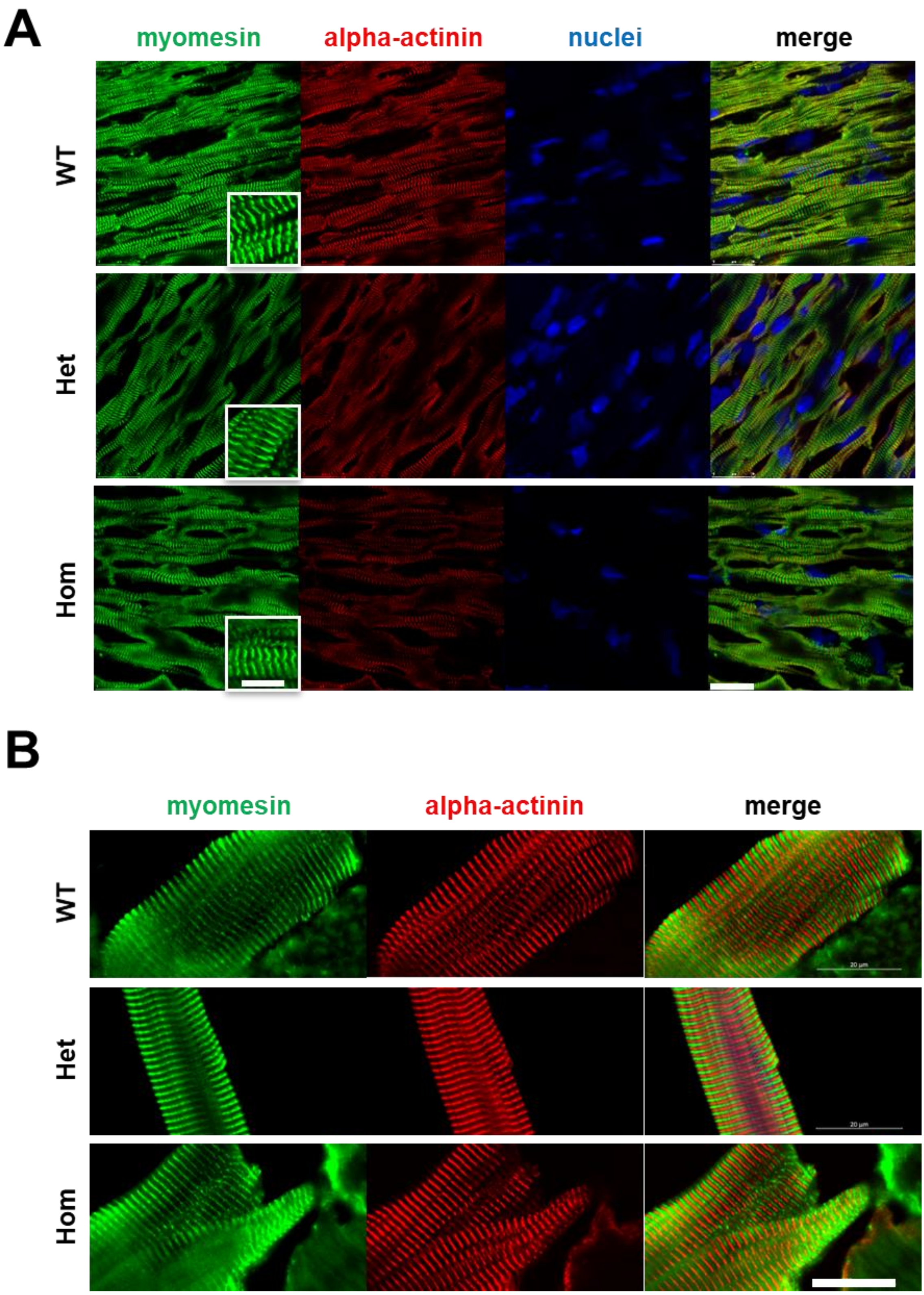
A
myomesin
alpha-actinin
nuclei
merge
WT
Het
Hom
B
myomesin
alpha-actinin
merge
WT
Het
Hom

**Figure S6. Isolated left ventricular cardiomyocytes from *Alpk3* K201X mice in the absence of the $Ca^{2+}$ indicator fura2 display hypercontractile unloaded sarcomere shortening with prolonged relaxation.** (**A**) Average sarcomere length traces from WT (blue), heterozygous (Het, red) and homozygous *Alpk3* K201X (Hom, grey) cardiomyocytes without fura2 loading. (**B**) Contractile traces following treatment with 0.5 μM mavacamten are shown in B. Dot plots for selected extracted parameters: basal sarcomere length (**C**), fractional shortening (**D**), T90% contraction (**E**) and T90% relaxation (**F**), respectively. Black lines represent median ± 95% confidence interval. Sample size (n) is representing cells per condition and is indicated. Statistical differences comparing WT to all other groups were calculated using a Kruskal-Wallis test with Dunn's test for multiple comparisons. All extracted parameters are tabulated in Supplementary Table S6C.

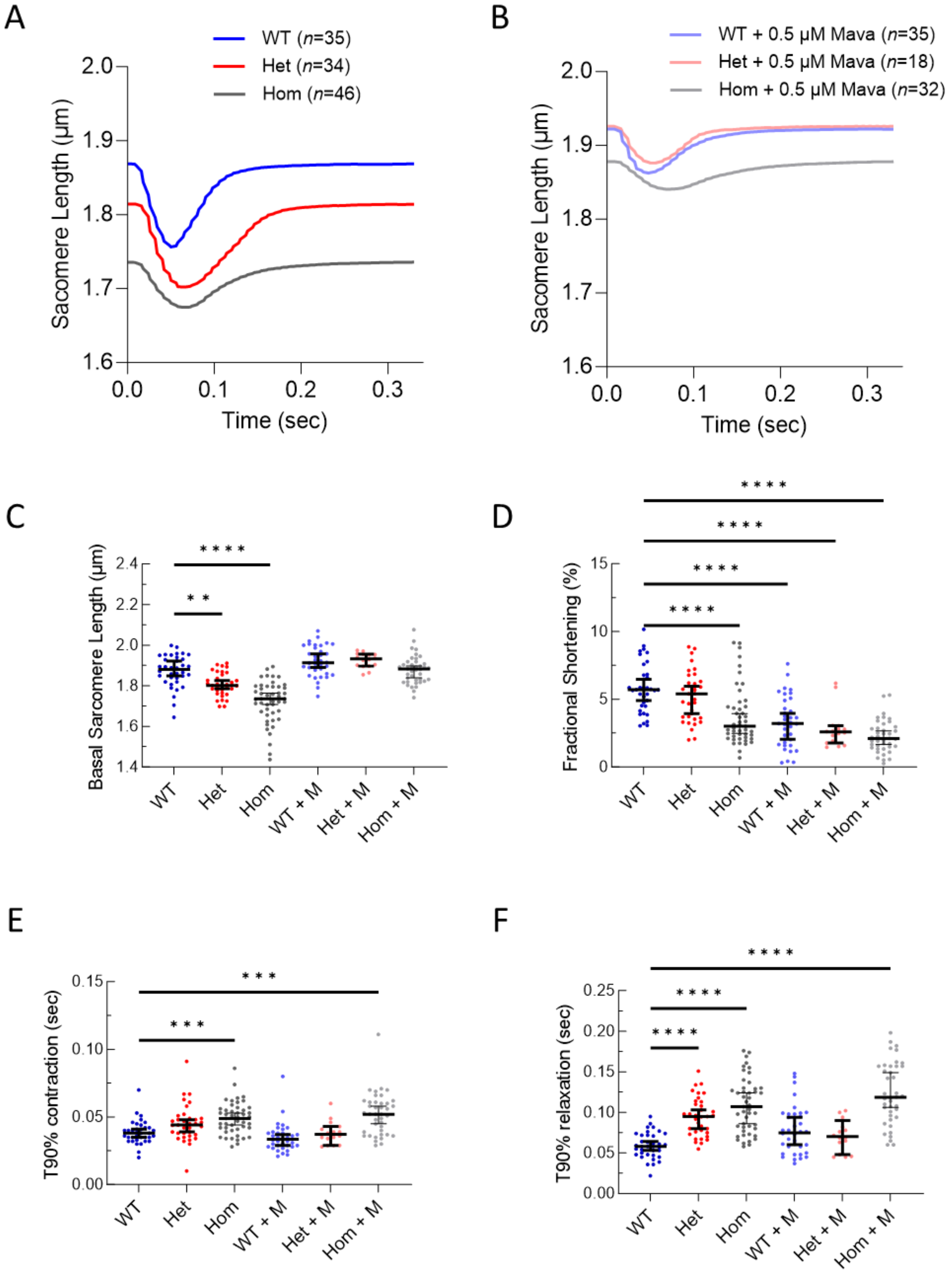
A
WT (n=35)
Het (n=34)
Hom (n=46)
Sacomere Length (µm)
Time (sec)
B
WT + 0.5 µM Mava (n=35)
Het + 0.5 µM Mava (n=18)
Hom + 0.5 µM Mava (n=32)
Sacomere Length (µm)
Time (sec)
C
Basal Sarcomere Length (µm)
WT
Het
Hom
WT + M
Het + M
Hom + M
D
Fractional Shortening (%)
WT
Het
Hom
WT + M
Het + M
Hom + M
E
T90% contraction (sec)
WT
Het
Hom
WT + M
Het + M
Hom + M
F
T90% relaxation (sec)
WT
Het
Hom
WT + M
Het + M
Hom + M

**Figure S7. Cluster analysis illustration from all significant extracted parameters taken from unloaded sarcomere shortening and calcium transient measurements.** Dots plots show parameters extracted from each cardiomyocyte split by individual mice for WT (blue), heterozygous (Het, red) and homozygous *Alpk3* K201X (Hom, grey) cardiomyocytes with and without treatment with 0.5 µM mavacamten. Parameters extracted include, basal sarcomere length (**A**), peak sarcomere length (**B**), fractional shortening (**C**), time to peak contraction (**D**), time to 50% contraction (**E**), time to 90% contraction (**F**), time to 50% relaxation (**G**), time to 90% relaxation (**H**), diastolic $[Ca^{2+}]_i$ (**I**), systolic $[Ca^{2+}]_i$ (**J**), $Ca^{2+}$ transient amplitude (**K**), time to peak $Ca^{2+}$ release (**L**), time to 90% $Ca^{2+}$ release (**M**), time to 10% $Ca^{2+}$ re-uptake (**N**), time to 50% $Ca^{2+}$ reuptake (**O**) and time to 90% $Ca^{2+}$ re-uptake (**P**). Black lines are the median error bars are 95% confidence interval. Statistical differences comparing WT to all other groups were calculated using nested 1-way ANOVA to adjust for hierarchical clustering of individual data sets between mice. P values in bold indicates significance p <0.05. Individual n numbers and all extracted parameters are tabulated in supplementary Table S6.

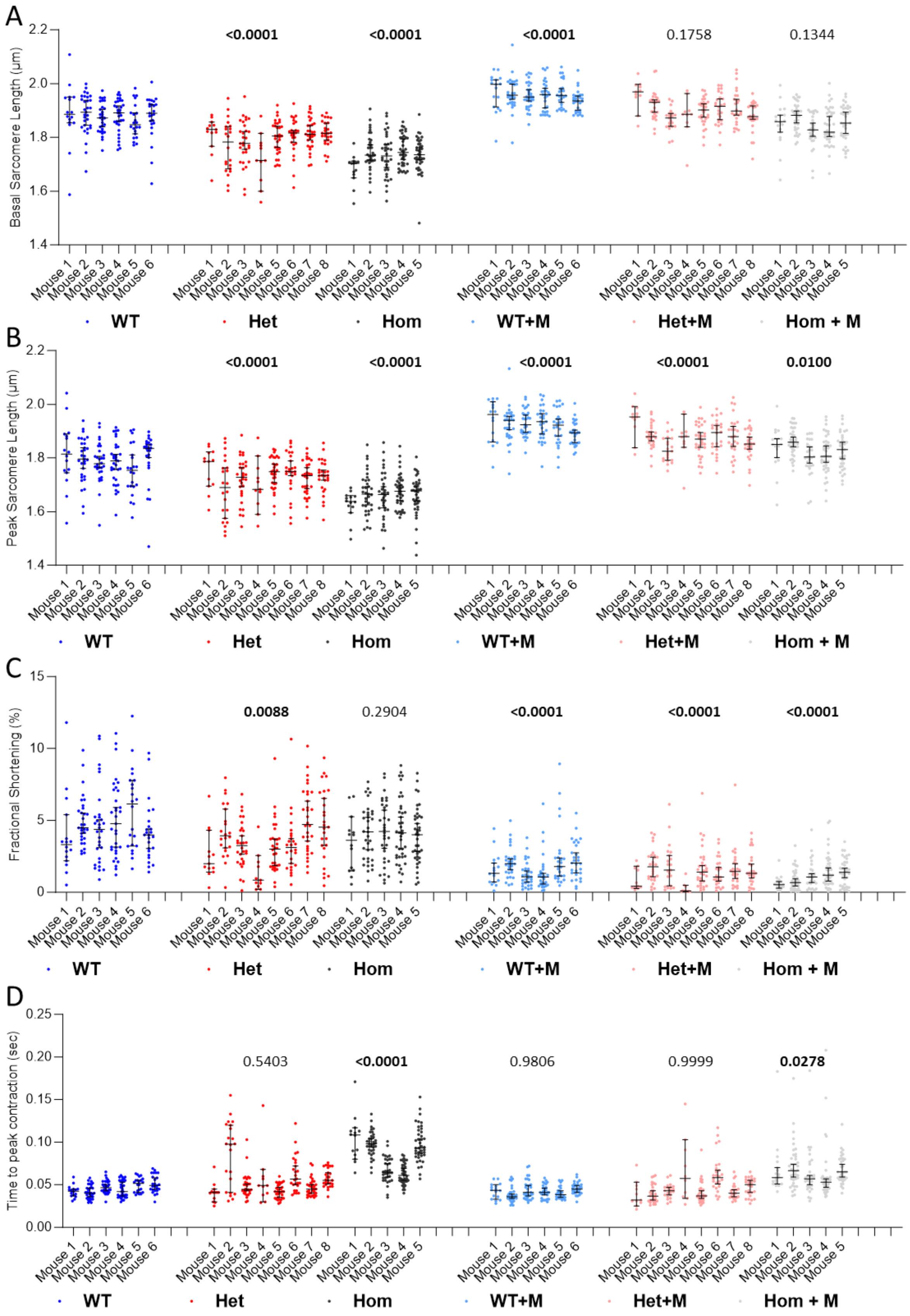
A
Basal Sarcomere Length (µm)
<0.0001
<0.0001
<0.0001
0.1758
0.1344
B
Peak Sarcomere Length (µm)
<0.0001
<0.0001
<0.0001
<0.0001
0.0100
C
Fractional Shortening (%)
0.0088
0.2904
<0.0001
<0.0001
<0.0001
D
Time to peak contraction (sec)
0.5403
<0.0001
0.9806
0.9999
0.0278
WT
Het
Hom
WT+M
Het+M
Hom + M

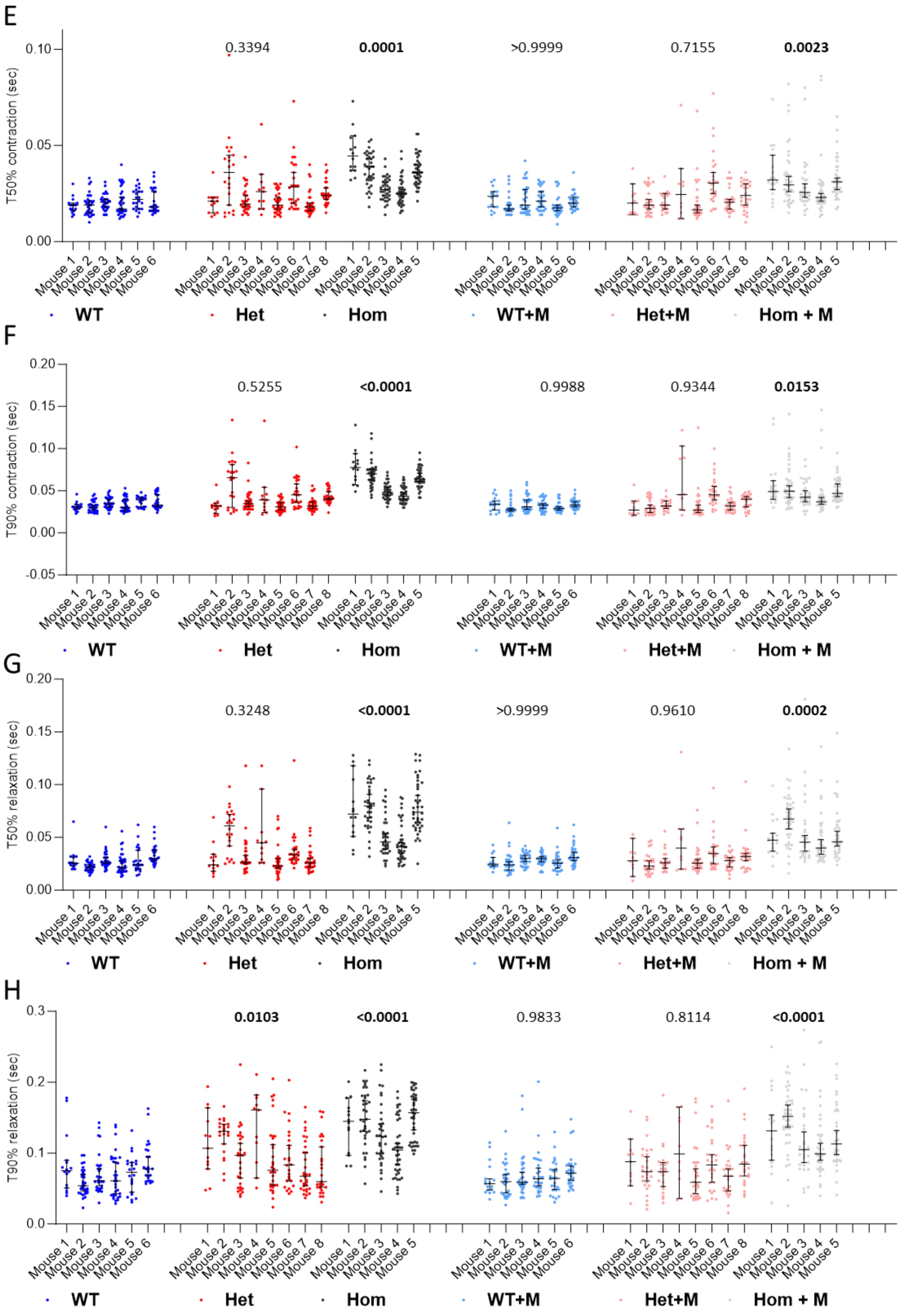
E
T50% contraction (sec)
0.10
0.05
0.00
0.3394
0.0001
>0.9999
0.7155
0.0023
Mouse 1 Mouse 2 Mouse 3 Mouse 4 Mouse 5 Mouse 6
Mouse 1 Mouse 2 Mouse 3 Mouse 4 Mouse 5 Mouse 6 Mouse 7 Mouse 8
WT Het Hom WT+M Het+M Hom + M
F
T90% contraction (sec)
0.20
0.15
0.10
0.05
0.00
-0.05
0.5255
<0.0001
0.9988
0.9344
0.0153
WT Het Hom WT+M Het+M Hom + M
G
T50% relaxation (sec)
0.20
0.15
0.10
0.05
0.00
0.3248
<0.0001
>0.9999
0.9610
0.0002
WT Het Hom WT+M Het+M Hom + M
H
T90% relaxation (sec)
0.3
0.2
0.1
0.0
0.0103
<0.0001
0.9833
0.8114
<0.0001
WT Het Hom WT+M Het+M Hom + M

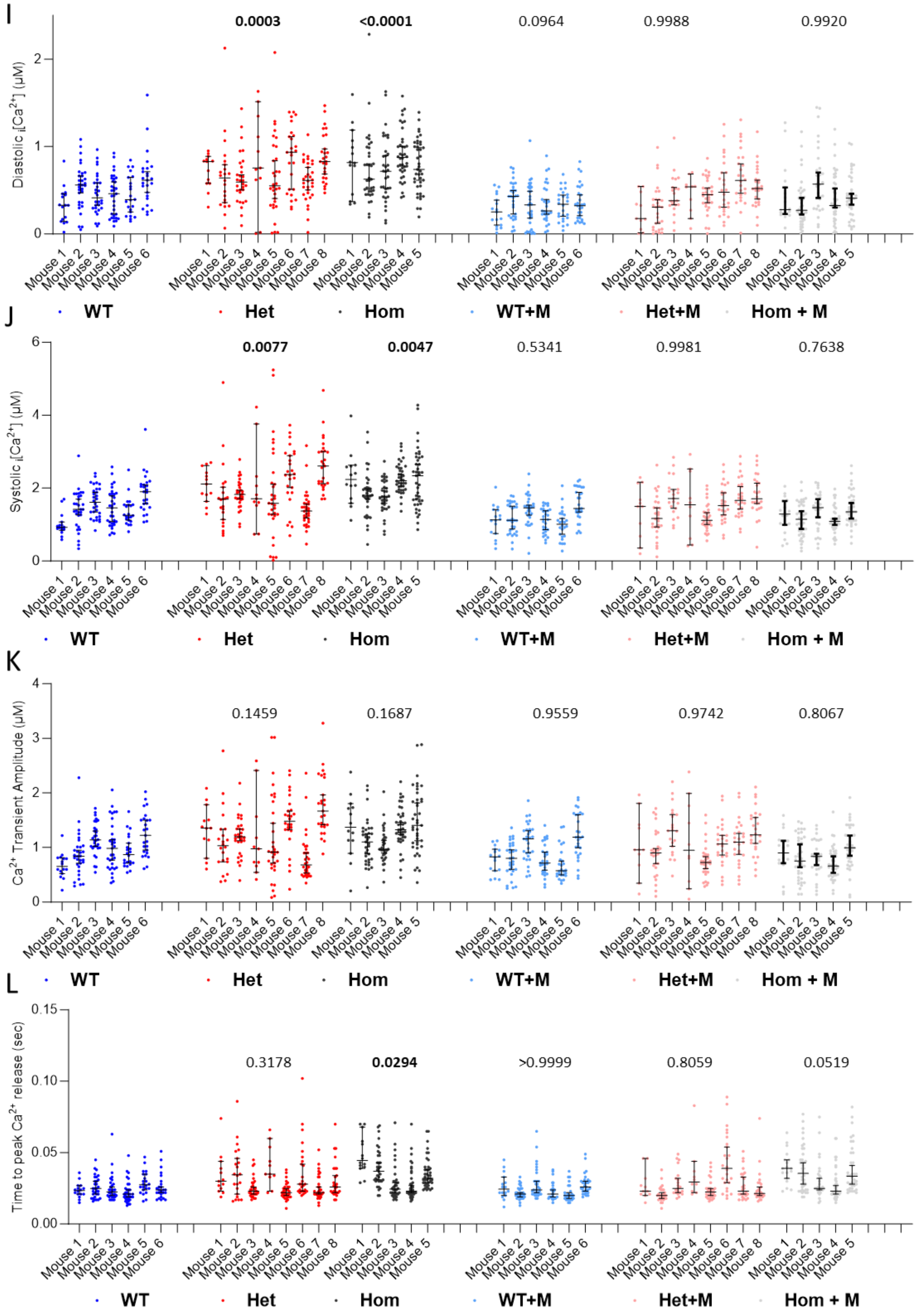
I
Diastolic $_i$[Ca$^{2+}$] (µM)
0.0003
<0.0001
0.0964
0.9988
0.9920
J
Systolic $_i$[Ca$^{2+}$] (µM)
0.0077
0.0047
0.5341
0.9981
0.7638
K
Ca$^{2+}$ Transient Amplitude (µM)
0.1459
0.1687
0.9559
0.9742
0.8067
L
Time to peak Ca$^{2+}$ release (sec)
0.3178
0.0294
>0.9999
0.8059
0.0519
Mouse 1
Mouse 2
Mouse 3
Mouse 4
Mouse 5
Mouse 6
Mouse 7
Mouse 8
WT
Het
Hom
WT+M
Het+M
Hom + M

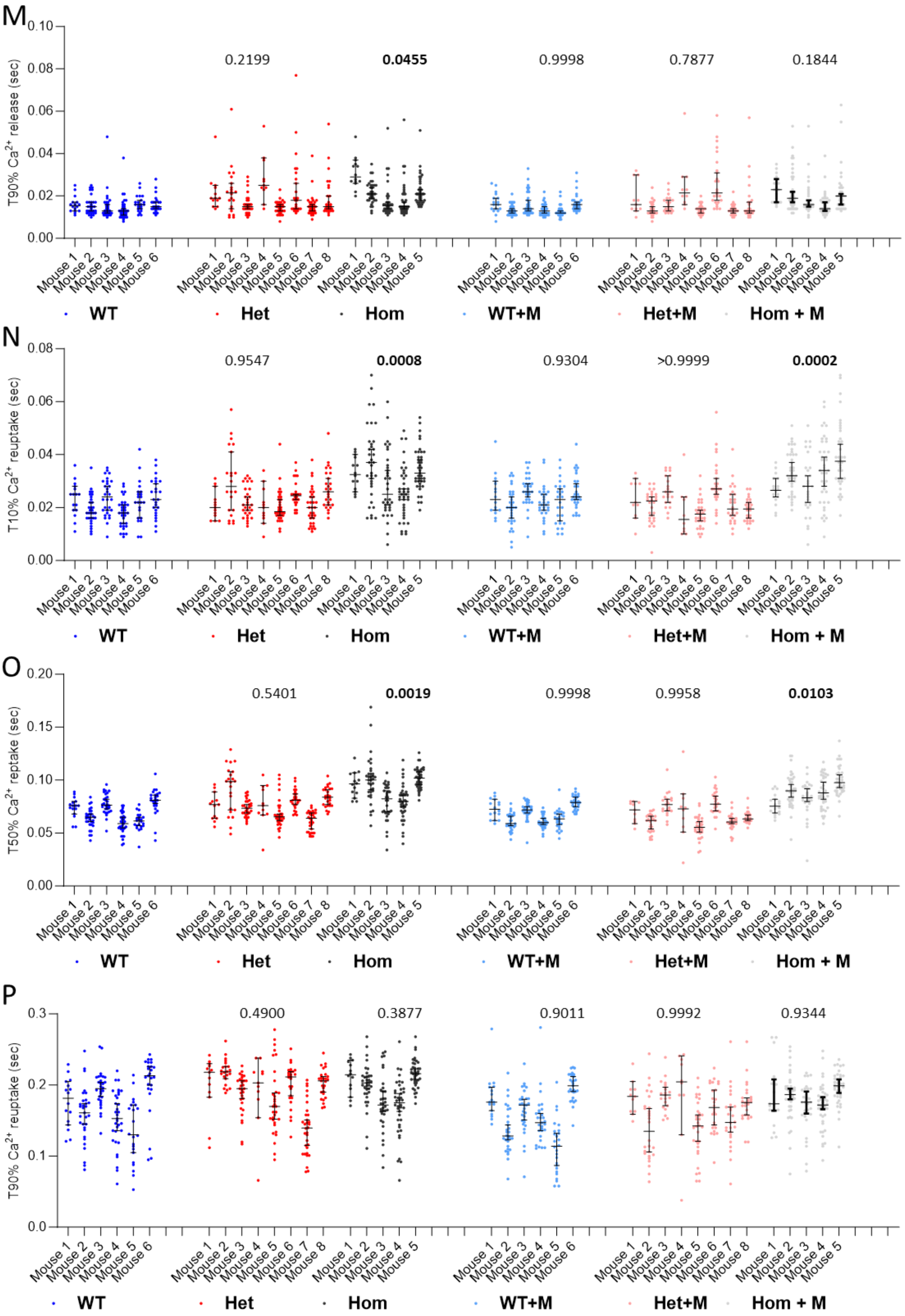
M
T90% Ca2+ release (sec)
0.2199
0.0455
0.9998
0.7877
0.1844
N
T10% Ca2+ reuptake (sec)
0.9547
0.0008
0.9304
>0.9999
0.0002
O
T50% Ca2+ reptake (sec)
0.5401
0.0019
0.9998
0.9958
0.0103
P
T90% Ca2+ reuptake (sec)
0.4900
0.3877
0.9011
0.9992
0.9344
Mouse 1
Mouse 2
Mouse 3
Mouse 4
Mouse 5
Mouse 6
Mouse 7
Mouse 8
WT
Het
Hom
WT+M
Het+M
Hom + M

**Figure S8. Whole western blot images.** Images show whole western blots that were zoomed (red boxes) to create **Figure 4A and C (blots in A and B respectively).** Red star in **B (top)** highlights overexposed / photobleached phospho-cTnI, red star in **B (bottom)** denotes BSA contaminant from cellular isolation. Both bands were manually removed from lane scan densitometry calculations.

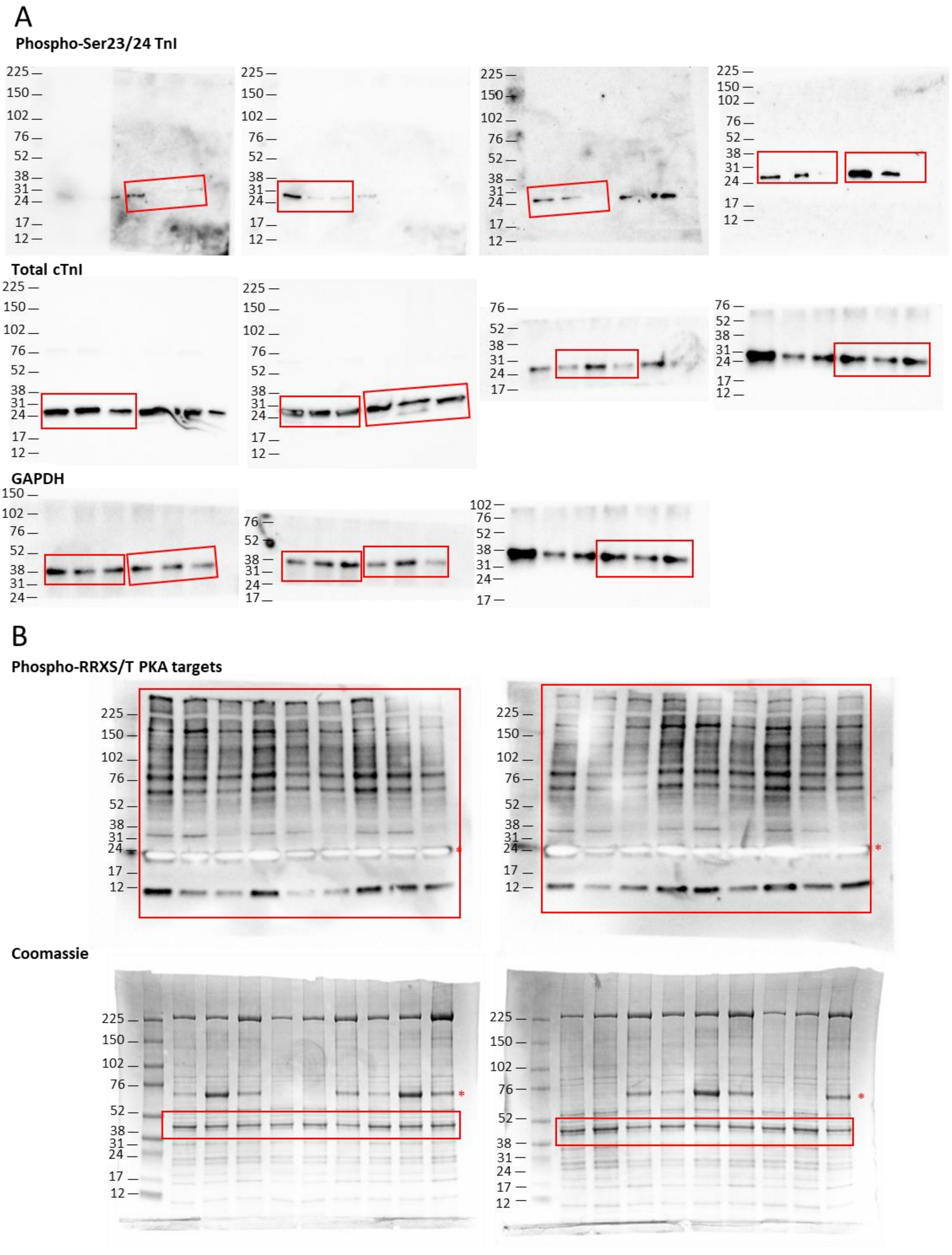

### References

[1] K. Gehmlich, M.S. Dodd, J.W. Allwood, M. Kelly, M. Bellahcene, H.V. Lad, A. Stockenhuber, C. Hooper, H. Ashrafian, C.S. Redwood, L. Carrier, W.B. Dunn, Changes in the cardiac metabolome caused by perhexiline treatment in a mouse model of hypertrophic cardiomyopathy, Mol Biosyst 11(2) (2015) 564-73.

[2] A.N. Abou Tayoun, T. Pesaran, M.T. DiStefano, A. Oza, H.L. Rehm, L.G. Biesecker, S.M. Harrison, G. ClinGen Sequence Variant Interpretation Working, Recommendations for interpreting the loss of function PVS1 ACMG/AMP variant criterion, Hum Mutat 39(11) (2018) 1517-1524.

[3] S. Richards, N. Aziz, S. Bale, D. Bick, S. Das, J. Gastier-Foster, W.W. Grody, M. Hegde, E. Lyon, E. Spector, K. Voelkerding, H.L. Rehm, A.L.Q.A. Committee, Standards and guidelines for the interpretation of sequence variants: a joint consensus recommendation of the American College of Medical Genetics and Genomics and the Association for Molecular Pathology, Genet Med 17(5) (2015) 405-24.
